# Genetic inactivation of the USP19 deubiquitinase regulates a-synuclein ubiquitination and inhibits accumulation of Lewy body like aggregates in mice

**DOI:** 10.1101/2022.12.21.521125

**Authors:** Lenka Schorova, Nathalie Bedard, Anouar Khayachi, Joao Bolivar-Pedroso, Hung-Hsiang Ho, Julie Huynh, Mikaela Piccirelli, Yifei Wang, Marie Plourde, Wen Luo, Esther del Cid-Pellitero, Irina Shlaifer, Yihong Ye, Thomas M. Durcan, Simon S. Wing

## Abstract

The USP19 deubiquitinase is found in a locus associated with Parkinson’s Disease (PD), interacts with heat shock proteins and promotes secretion of a-synuclein (a-syn) through the misfolding associated protein secretion (MAPS) pathway. Since these processes might modulate the processing of a-syn aggregates during the progression of PD, we tested the effect of USP19 knockout (KO) in mice expressing the A53T mutation of a-syn and in whom a-syn preformed fibrils (PFF) had been injected in the striatum. Compared to WT, KO brains showed decreased accumulation of phospho-synuclein (pSyn) positive aggregates. The improved pathology was associated with less activation of microglia, higher levels of synaptic marker proteins and improved performance in a tail suspension test. Exposure of primary neurons from WT and KO mice to PFF in vitro also led to decreased accumulation of pSyn aggregates. KO did not affect uptake of PFF in the cultured neurons. It also did not affect the propagation of aggregates as assessed by exposing WT or KO neurons to PFF and measuring pSyn positive aggregates in non-exposed adjacent neurons separated using a microfluidics device. We conclude that USP19 instead modulates intracellular dynamics of aggregates. Indeed, at the early time following PFF injection when the number of pSyn positive neurons were similar in WT and KO brains, the KO neurons contained less aggregates. KO brain aggregates stained more intensely with anti-ubiquitin antibodies. Immunoprecipitation of soluble proteins from primary neurons exposed to PFF with antibodies to ubiquitin or pSyn showed higher levels of ubiquitinated a-syn oligomeric species in the KO neurons. We propose that the improved pathology in USP19 KO brains may arise from decreased formation or enhanced clearance of the more ubiquitinated aggregates and/or enhanced disassembly towards more soluble oligomeric species. USP19 inhibition may represent a novel therapeutic approach that targets the intracellular dynamics of a-syn complexes.

## INTRODUCTION

Parkinson’s disease (PD) is the second most common neurodegenerative disease and is characterized by motor and non-motor dysfunctions due to the progressive loss of dopaminergic neurons. Current therapies target primarily symptoms and are often hampered by adverse effects. With age as the greatest risk factor and the predicted growth in the elderly population, the prevalence of PD is expected to increase rapidly. Therefore, it is imperative to define pathogenetic mechanisms to identify new therapeutic targets.

A pathological hallmark of PD is the hierarchical and progressive spread of intracellular misfolded protein inclusions, termed Lewy bodies (LBs), between interconnected brain regions^1,2^. One of the major components of LBs is a-synuclein (a-syn), an abundant brain protein accumulated in both sporadic and familial forms of PD as well as in other related disorders collectively termed synucleopathies. Under normal conditions, a-syn predominantly localizes in axons where it regulates presynaptic vesicle trafficking and neurotransmitter release by acting as a SNARE complex chaperone^3^. A-syn is an intrinsically unstructured protein natively occurring as monomers which are prone to misfolding and fibril formation^4^. Mutations, posttranslational modifications as well as association with chaperones can modulate its propensity to aggregate^5-7^. Indeed, several a-syn missense mutations (A30P^8^, E46K^9^, A53T^10^, A53E^11^, H50Q^12^ and G51D^13^) as well as duplication^14^ and triplication^15^ of the a-syn gene SNCA all cause familial PD. Importantly, a-syn-containing aggregates can propagate between brain cells, which was initially suggested by the observation that grafted fetal mesencephalic neurons stained positively for a-syn pathology years after transplantation into the brain of PD patients^16,17^. Since then, several studies confirmed the transmission of a-syn aggregates in animal models^18-20^. Moreover, transmitted a-syn can act as seeds converting monomeric a-syn in recipient cells into toxic oligomers or fibrils, thus amplifying a-syn aggregation-associated cytotoxicity in neurons^21-23^. Thus, pathways that actively participate in the transmission of aggregated a-syn are of interest as targets for treatment of PD and other diseases associated with protein misfolding and prion-like spread. To date, two mechanisms have been proposed to mediate the intercellular spreading of a-syn aggregates: intercellular nanotubes or secretion of a-syn by unconventional pathways followed by its internalization by naïve neurons.

The internalization of fibrillar a-syn takes place, at least in part, via dynamin-dependent endocytosis^24^. Studies have reported different cell surface candidates that recruit a-syn to the plasma membrane to mediate its uptake. Heparan sulfate proteoglycans (HSPGs) comprise of several plasma membrane and extracellular glycoproteins that can mediate the recruitment and internalization of many cargos including fibrillar a-syn but not soluble oligomers in both neuronal-like cells^25,26^ and in primary neurons^27^. Lymphocyte-activation gene 3 (LAG3) also mediates the endocytosis of a-syn fibrils. Genetic depletion or antibody attenuation of LAG3 reduced a-syn fibril endocytosis, neuron-to-neuron spread and delayed the a-syn preformed fibril (PFF)-induced neuronal loss in the mouse substantia nigra^28^. A-syn is constitutively acetylated at the N terminus which appears to be important for binding to membrane N-glycans in primary neurons and neuroblastoma cells but not HEK cells. The cleavage of N-glycans reduces the internalization of a-syn monomers and PFF. In addition to LAG3, this study also demonstrated that the depletion of the membrane glycoprotein neurexin 1β blocked the entry of acetylated a-syn monomers and PFF^29^. A recent study in U2OS and iPSC-derived human dopaminergic neurons and astrocytes reported that a-syn PFF are internalized via macropinocytosis and trafficked to lysosomes and multivesicular bodies within minutes of exposure. These findings put in question previous reports that are in favor of a-syn internalization via dynamin dependent endocytosis^30^.

Although significant insights have been obtained on a-syn uptake, much less is known about how it is released from cells. A-syn lacks a secretory sequence and therefore is not secreted via the conventional ER-Golgi pathway. Several lines of evidence suggest that a-syn can be secreted via non-canonical vesicle-mediated exocytosis. While a small fraction of extracellular a-syn appears to be released together with exosomes^30,31^, other studies suggested an exosome independent unconventional protein secretion pathway termed as misfolding-associated protein secretion (MAPS) to specifically target misfolded proteins including a-syn and tau for secretion^32-34^. MAPS can be initiated when misfolded proteins are recruited by the endoplasmic reticulum (ER)-embedded deubiquitinase USP19. Subsequently, cargo proteins are handled by HSP70 and its co-chaperone DNAJC5, before they enter the lumen of a peri-nuclear membrane compartment and late endosomes^35^. Cargos are further directed to the cell exterior or to lysosomes for degradation.

In addition to protein secretion and lysosomal degradation, USP19 has also been implicated in degradation of two major autophagy players Beclin 1^36^ and TBK1^37^. Given the reported association of USP19 with two major cytosolic chaperones HSC70 and HSP90^38,39^, it is anticipated to function as a key ubiquitin processing enzyme in protein homeostasis regulation. In view of these functions of USP19 in proteostasis and the recent recognition that it is located in a locus on chromosome 3 that is associated with Parkinson’s disease^40^,(https://pdgenetics.shinyapps.io/GWASBrowser/), we tested whether USP19 regulates a-syn processing and/or propagation in PD pathology. We used USP19 KO mice to study the effects of USP19 depletion on PD-like pathology *in vivo* and in primary neurons. We report that USP19 depletion significantly reduces pSyn pathology in a mouse model expressing the human a-syn A53T disease-causing mutation as well as in primary neurons and this effect is primarily due to altered intracellular handling of pathological species of a-syn.

## RESULTS

### USP19 is expressed in the adult mouse brain

USP19 is expressed as two major isoforms – one cytoplasmic and the other ER localized - which arise from alternative splicing of the final exon that encodes a transmembrane domain. As antibodies that recognize the small differences in amino acid sequence are not available, we first characterized the expression and localization of USP19 mRNA in mouse brains by in situ hybridization with USP19 oligo probes specifically recognizing mRNAs of different USP19 isoforms. The USP19 ER isoform (USP19-ER), which specifically mediates the MAPS pathway, was detected across multiple regions of the USP19 WT but not the KO brains (Fig. 1A, B). Specifically, USP19-ER mRNA was present in neurons (identified as NeuN positive) including dopaminergic tyrosine hydroxylase (TH)-positive neurons (Fig. 1C). Additionally, USP19-ER mRNA was also detected in astrocytes and microglia (Fig. 1C). Similarly, oligo probes targeting specifically the USP19 cytoplasmic isoform (USP19-cyt) and well as both isoforms (USP19-common) mRNA confirmed that the cytosolic isoform is also expressed in the mouse brain (Fig. S1A, B). Immunoblotting of protein from whole brain homogenates confirmed the expression of USP19 protein in WT and its absence in KO animals (Fig. 1D). These data confirm the expression of USP19 in the mouse brain and suggest a possible role for USP19 in protein homeostasis in various cell types in this tissue.

**Fig 1:**
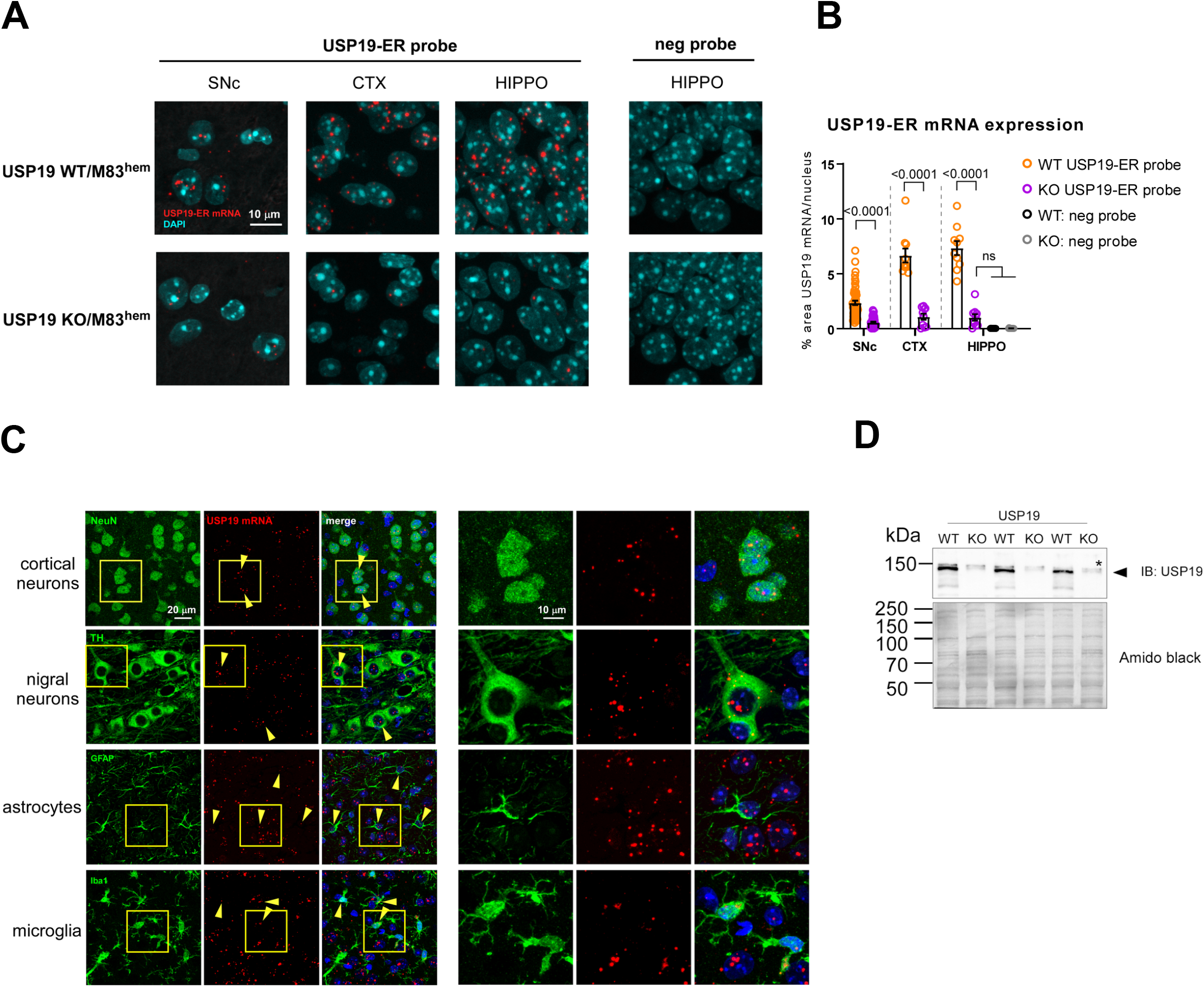
USP19-ER expression in the mouse brain. **A**. RNAScope in-situ hybridization on formalin-fixed paraformaldehyde-embedded (FFPE) brain sections to detect the presence (red signal) of ER-specific USP19 mRNA isoform in USP19 WT and its absence in KO/M83^hem^ animals. Shown are substantia nigra pars compacta (SNc), cortex (CTX) and hippocampus (HIPPO). A probe targeting a bacterial mRNA was used as a negative control probe. DAPI was used to label nuclei. **B**. Quantification of RNAScope USP19-ER signal presented as percent area of red pixels in >40 TH-positive cells in SNc, and 10 randomly selected nuclei in CTX and HIPPO. **C**. RNAScope FISH coupled with IF of neuronal (NeuN, TH) and glia markers (Iba1, GFAP) showing USP19-ER mRNA expression in multiple brain cell types. **D**. Immunoblot of whole-brain lysates showing the expression of USP19 in USP19-WT and -KO/M83^hem^ brains. Asterisk highlights a non-specific band, arrowhead points to full length USP19. Data are mean ± s.e.m. Unpaired t-test and one-way ANOVA were used for statistical analysis followed by Tukey’s multiple comparisons test. P-values are indicated.

### Loss of USP19 improves pSyn and pTau pathology in a PD-like mouse brain

To explore the involvement of USP19 in PD pathology, we crossed USP19 WT and KO mice to a transgenic line expressing the human PD-causing mutant a-syn^A53T^ (M83 hemizygous transgene [M83^hem^]) under the CNS prion promoter^41^. M83^hem^ mice spontaneously develop age-dependent motor impairment leading to paralysis between 22 and 28 months of age, but the propagation of ectopically expressed a-syn is not obvious^41^. To accelerate the development of a-syn propagation and the associated PD-like pathology in a robust manner in a mouse model, we injected preformed a-syn fibrils (PFF) or control a-syn monomers in the dorsal striatum at 3 months of age, which typically leads to Lewy body-like pathology and motor defects within 3-4 months^42^. At the onset of motor symptoms (90-110 days post-injection [dpi]), mice were sacrificed, and the brains analyzed by immunohistochemistry. To detect Lewy body-like pathology, we used a validated phosphoS129-syn (pSyn) antibody (Fig. S2A) to stain coronal brain sections, which revealed widespread accumulation of pSyn, a hallmark of Lewy body pathology in PD, in WT mice injected with a-syn PFF. Consistent with previous studies^43^, no pSyn-containing cells were found in KO or WT animals that had been injected with a-syn monomers (Fig. 2A, B). In general, more pSyn signals were detected on the ipsilateral side (side of injection) than the contralateral side, suggesting that the Lewy body-like pathology occurs first in the injected region and later spreads to the contralateral side of the brain. Interestingly, fewer pSyn-positive (pSyn^+^) cells were detected in USP19 KO mice exposed to PFF in many of the ipsilateral brain regions including the substantia nigra pars compacta (SNc), striatum (Str), periaqueductal grey (PAG), piriform cortex (Piri; Fig. 2A, B), cortex (Ctx), hippocampus (Hippo), hypothalamus (Hypo), midbrain, olfactory bulb (OB) and thalamus (Thal; Fig. S2B). Similarly, the number of pSyn^+^ cells was significantly reduced in the KO contralateral PAG, Piri (Fig. 2A, B), and Hypo (Fig. S2B) compared to WT counterparts. Integrating the results from all the individual regions of the brain showed that although some ipsilateral regions were more affected by USP19 KO than their contralateral counterparts (SNc, Str and PAG) (Fig. S2C), an overall significant reduction in pSyn pathology was found in KO ipsi- and contralateral regions compared to the counterpart regions in WT animals (Fig. 2C), suggesting a less efficient progression of the PD-like pathology in the brains of USP19 KO animals than in WT brains. Interestingly, we observed higher levels of pSyn pathology in USP19 WT females relative to WT males (Fig. S2F, G) which could be due to the positive regulation of USP19 expression by estrogen^44^.

**Fig. 2:**
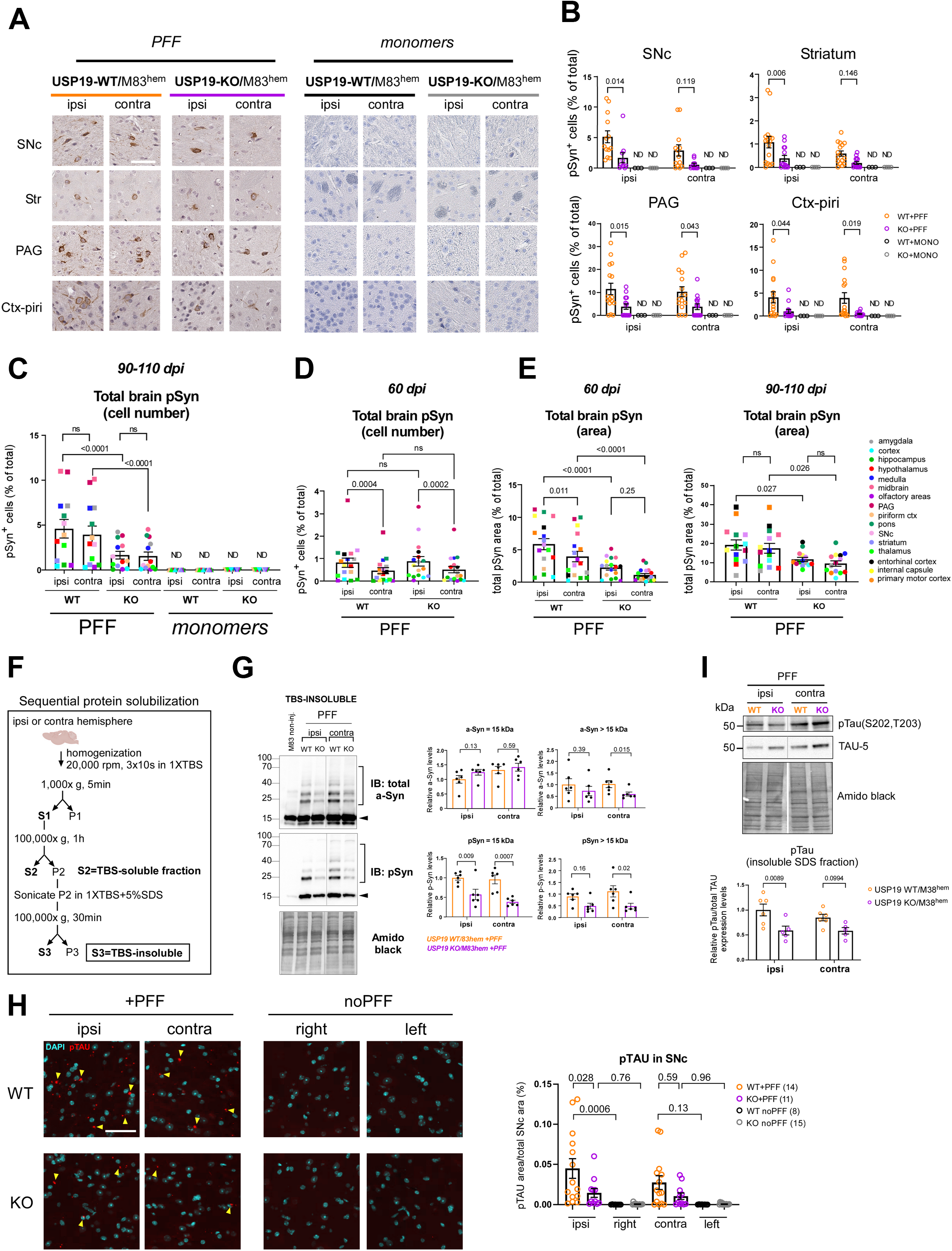
USP19 depletion improves pSyn and pTAU pathology in a mouse model of Parkinson’s disease. **A**. Representative photomicrographs of IHC staining of pS129-Syn (brown) from coronal brain sections of substantia nigra pars compacta (SNpc), striatum (Str), hippocampus (Hippo) and piriform cortex (Piri) from PFF- or a-syn monomers-injected USP19-WT or KO/M83^hem^ mice at 90-110 dpi (days post injection). Scale bar, 50 µ m. Sections are counterstained for the presence of nuclei with hematoxylin (purple). Ipsi (ipsilateral) indicates the injected hemisphere, contra (contralateral) indicates the non-injected hemisphere. **B**. Quantification of pS129-Syn^+^ cells using QuPath analysis in indicated brain regions. Each data point represents a mean of 4 brain sections per animal. n ≥ 10 (PFF-injected) and n=3-4 (a-syn monomers-injected) biologically independent animals. Data are mean ± s.e.m. Two-way ANOVA was used for statistical analyses followed by Sidak multiple comparisons test. **C and D**. Quantification of total brain pS129-Syn represented as pSyn^+^ cells (% of total) using IHC in B and C at 90-110 dpi (C) and 60 dpi (D). Each point is a mean of means per brain region (each brain region is color-coded). Data are mean ± s.e.m. Two-way ANOVA was used for statistical analysis followed by Tukey’s multiple comparisons test **E**. Overall quantification of total brain pS129-Syn represented as total pSyn area (% of total) using IF at 60 dpi and 90-110 dpi. Each point is a mean of means per brain region. Data are mean ± s.e.m. Two-way ANOVA was used for statistical analysis followed by Tukey’s multiple comparisons test. **F**. Scheme of sequential protein solubilization protocol used to generate data in 2F, G and S2 F, G. **G**. Representative immunoblots of total a-syn and pS129-Syn of insoluble fractions (SDS-solubilized) of the whole brain hemispheres. Arrowhead points to monomeric a-syn or pS129-Syn. Square bracket highlights oligomeric species of a-syn or pS129-Syn. Amido black staining was used as a loading control. Quantification graphs of a-syn and pS129-Syn monomer and oligomers are shown separately. Data are mean ±s.e.m., n=6 biologically independent animals. Unpaired Mann-Whitney t-test was used for statistical analysis. **H**. Representative images of IF staining of pTAU in SNc of PFF-injected mice and noninjected controls (noPFF) at 90-110 dpi. n ≥ 8 biologically independent animals. Data are mean ±s.e.m. One-way ANOVA was used for statistical analysis followed by Tukey’s multiple comparisons test. **I**. Representative immunoblots of total tau (TAU-5) and p-Tau (S202, T203) of insoluble fractions (SDS-solubilized) of the whole brain hemispheres. Amido black staining was used as a loading control. Quantification of pTau was normalized to total tau of 5-6 independent animals per genotype. Data are mean ±s.e.m. Two-way ANOVA with Sidak multiple comparisons test was used for statistical analysis.

To begin exploring whether the reduction in pSyn was due to slower propagation or differential formation/clearance of a-syn aggregates over time in the KO mice, we performed the same analysis at an earlier time point post-PFF injection. At 60 dpi, there was no difference in the number of pSyn^+^ cells between WT and KO animals suggesting that pSyn pathology develops initially in similar number of cells and therefore USP19 may not be involved in the initial cell-to-cell propagation mechanism (Fig. 2D). As pSyn^+^ aggregates localize not only to the cell body (Lewy body-like structures) but also to neurites (Lewy neurite-like structures), we analyzed the total area of pSyn^+^ staining in several brain regions between WT and KO animals at both 60 and 90-110 dpi. A significantly smaller pSyn^+^ area was found in both ipsi- and contralateral regions of KO compared to WT animals (Fig. 2E). Interestingly, the reduction in pSyn^+^ area is already apparent at 60 dpi at a time when the number of pSyn^+^ cells are similar. These data indicate that the loss of USP19 decreases the intraneuronal accumulation of pSyn pathology.

To confirm and extend these findings, we quantified by immunoblotting, pSyn and total a-syn in TBS soluble (Tris-Buffed Saline-solubilized; Fig. S2D, E) and TBS insoluble fractions (TBS-insoluble, SDS-solubilized) of homogenates prepared from ipsi- and contralateral hemispheres (Fig. 2F, G) as the presence of insoluble pSyn is another hallmark of Lewy body pathology. There was significantly less TBS-insoluble pSyn (15-kDa band representing SDS-solubilized pSyn) in both ipsi and contralateral KO brains when compared to WT brains. A significant reduction in high MW pSyn species (>15 kDa, reflecting likely posttranslationally-modified pSyn or complexes resistant to SDS dissociation) was also seen in the KO brain. Additionally, we detected a significant decrease in insoluble high MW total a-syn species in contra and a similar trend in ipsi KO brains (Fig. 2G). By contrast, total TBS-insoluble monomeric a-syn (15-kDa band representing SDS-solubilized a-syn) was largely unchanged between genotypes. There was no difference between WT and KO in TBS-*soluble* total a-syn and pSyn (Fig. S2D, E). Thus, there were decreased levels of specifically pathological forms of a-syn (phosphorylated and insoluble) in the KO brains.

Cooccurrence of pathological accumulation of a-syn and tau has been reported in AD and PD suggesting these proteins may synergistically interact to promote the accumulation of each other^45-48^. Therefore, we hypothesized that USP19 could also regulate the level of tau pathology in the M83 mouse model which develops tau inclusions^49^. To analyze the degree of tau pathology, we first validated the specificity of the pTau antibody (Fig. S2A) and then quantified pTau levels in SNc by immunofluorescence (IF). Interestingly, pTau inclusions were significantly reduced in ipsi-SNc with a similar trend in contralateral SNc of KO mice (Fig. 2H). Western blot analysis corroborated our imaging data and revealed a significant reduction of total brain insoluble pTau in ipsilateral hemisphere with a less evident difference in the contralateral hemispheres in KO animals (Fig. 2I). Thus, USP19 regulates both a-syn and tau aggregate accumulation.

### Loss of USP19 reduces PD-pathology-associated neuroinflammation in mice

Neuroinflammation is a prominent hallmark of PD and many rodent PD models and a key contributor to the pathogenesis of the disease^50-53^. In response to various insults, including the administration of PFF, microglia undergo deramification and exhibit an amoeboid morphology during the neuroinflammatory response^54-56^. To determine whether the loss of USP19 results in changes of microglia morphology post-PFF administration, we measured the mean size of Iba1-positive (Iba^+^, marker of microglia) cells by immunostaining. We found that microglia were significantly smaller in the ipsilateral midbrains of KO compared to WT animals, with a trend to a similar reduction in the contralateral midbrains (Fig. 3A, B). Infiltration of microglial cells to the site of inflammation to combat infection and phagocytose cell debris is another indicator of increased neuroinflammation^54-56^, which can be determined by measuring the density of Iba+ staining. Therefore, we assessed microglial density by measuring total Iba^+^ area. There was a ∼50% decrease in Iba1^+^ cell density in KO ipsi- and contralateral midbrain compared to WT brains (Fig. 3A, B). Moreover, analysis of additional brain regions revealed a decrease in both ipsi- and contralateral microglia size and density in KO+PFF mice (Fig. 3C, 3D). In further support of this, immunoblot analysis revealed decreased astrocytic (GFAP) and microglia (Iba1) markers in total brain homogenates of KO animals (Fig. 3E, F). Taken together, the loss of USP19 leads to a decrease in the neuroinflammatory response which is consistent with the reduced pSyn pathology in the KO animals.

**Fig. 3:**
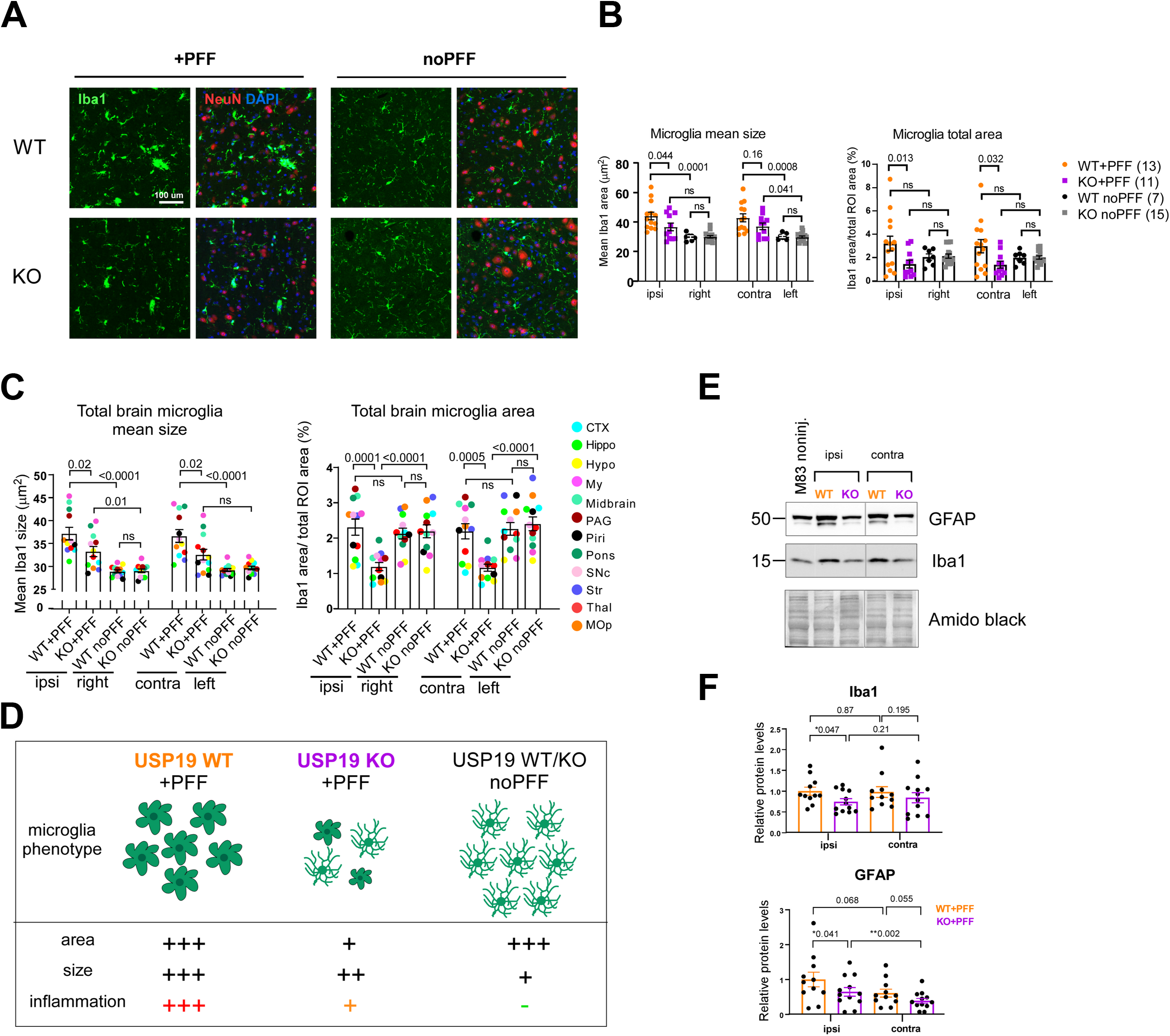
USP19 depletion reduces markers of neuroinflammation in a mouse model of Parkinson’s disease. **A**. Representative photomicrographs of IF staining for the microglia marker Iba1 (green) and the neuronal marker NeuN (red) on brain sections. DAPI was used to label nuclei. PFF-noninjected right and PFF-injected ipsilateral hemispheres of the midbrain region are shown. Scale bar, 100 μm. **B**. Quantification of the mean size of Iba1^+^ cells and Iba1 density (represented as total Iba1 area over total area of midbrain). n≥ 7 biologically independent animals. Two-way ANOVA with Tukey’s multiple comparison test was used for statistical analysis. **C**. Overall quantification of Iba1 IF staining in 12 brain regions on FFPE sections. The mean size (C) and Iba1 density (D), represented as total Iba1 area over total area of region of interest, are plotted. Each brain region is color-labeled. Each data point is a mean of means per brain region. n≥ 7 biologically independent animals. Two-way ANOVA with Tukey’s multiple comparison test was used for statistical analysis. P-values are indicated. **D**. Schematic summarizing microglia phenotype in PFF-treated and -nontreated USP19 WT and KO/M83^hem^ mice. PFF triggers microglia activation (gliosis) in WT and to a lesser degree in KO animals. Moreover, KO animals show reduced microglia population. **E. and F**. Representative immunoblots (E) and quantification (F) of glial markers known to be upregulated in neuroinflammation: GFAP (astrocytes) and Iba1 (microglia) normalized to amido black staining. PFF-injected (ipsi) and non-injected (contra) whole hemispheres were used. n ≥ 10 biologically independent animals. Data are mean ±s.e.m. Unpaired t-test was used for statistical analysis. P-values are indicated.

### Loss of USP19 improves synaptic markers and behavioral outcome in PD-like mice

The overexpression of M83 transgene and intrastriatal PFF inoculation has been previously shown to trigger a progressive loss of dopaminergic neurons in the substantia nigra characterized by lower levels of tyrosine hydroxylase (TH), an enzyme involved in dopamine production^42,43^. As the loss of USP19 led to a decrease in pSyn and pTau pathology and reduced neuroinflammation in PD-like mice, we hypothesized that these changes may result in a neuroprotective phenotype and lead to decreased neurodegeneration in KO animals. To test this, we measured markers of dopaminergic neurons. Immunoblot analysis of whole ipsi- and contralateral brain homogenates revealed in KO+PFF brains, a trend to higher protein levels in ipsilateral TH, a significant increase in ipsilateral synaptotagmin-1, a presynaptic calcium sensor, and contralateral SNAP25, a presynaptic vesicle marker and SNARE complex member (Fig. 4A, B). The higher protein levels in the KO brain indicate that USP19 loss may partially protect against neurodegeneration.

**Fig. 4:**
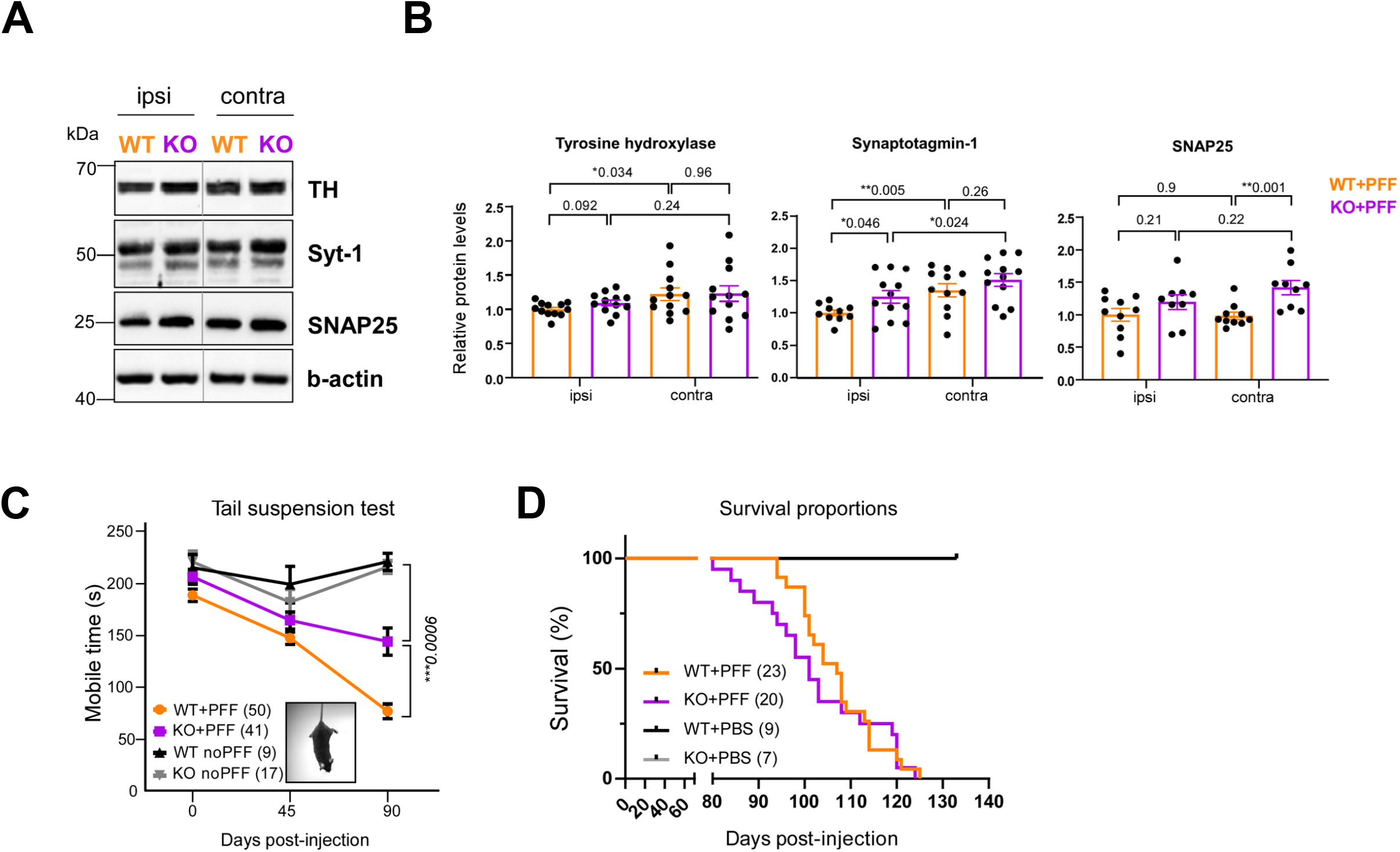
Loss of USP19 improves synaptic markers and behavioral outcome in PD-like mice. **A. and B**. Representative immunoblots (C) and quantifications (D) of synaptic markers TH (dopaminergic synapse), synaptotagmin-1 and SNAP25 (excitatory presynapse) using whole ipsi- and contralateral brain homogenates. n ≥ 9 biologically independent animals. Data are mean ±s.e.m. Unpaired t-test was used for statistical analysis. **C**. Tail-suspension test as a measure of anxiety-like behaviors and mobility, represented as a total mobile time (sec) defined as time of a mouse actively trying to get to the upright position in oppose to passive hanging. TST was performed at three time points: pre-PFF, 1.5 and 3-months post-PFF. Non-injected age-matched animals were used as controls. Two-way ANOVA with Tukey’s multiple comparison test was used for statistical analysis. **D**. Kaplan-Meier survival analysis of USP19 WT/M83^hem^ and KO/M83^hem^ animals injected with PFF or PBS. n≥ 7 biologically independent animals. Statistical analysis for survival curves was performed by long-rank (Mantel-Cox) test. P-values are indicated.

As previously shown, M83 and PFF-inoculated mice develop motor and cognitive function deficits related to PD^43,52,57^. To assess the effect of USP19 loss on both motor and cognitive functions in PD-like mice, we performed behavioral phenotyping. Although no differences between WT and KO mice injected with PFF were detected in wire hang, grip strength or rotarod tests (data not shown), KO animals showed a significant improvement in mobile time in the tail suspension test (TST). The absence of mobility in TST relates to the inability to get to the upright position which can reflect a defect in motor function (Fig. 4C). Finally, a survival experiment showed no improvement in overall survival rate in USP19 KO mice. In addition, no differences in survival were reported between sexes (Fig. 4D, S3A). However, KO mice were smaller in weight (Fig S3B) which may be a confounding factor that impairs survival under stress conditions. Taken together with the pathological findings described above, the loss of USP19 leads to a favorable brain phenotype in PFF-injected mice, but no apparent beneficial effect on overall survival.

### The loss of USP19 in neuronal cells reduces pSyn^+^ fibrils without affecting uptake of fibrils or propagation to other neurons

We next used primary neurons to explore the potential mechanisms underlying USP19’s neuronal protective role against pSyn accumulation. PFF treatment of primary rodent neurons leads to a progressive accumulation of pSyn^+^ fibrils resulting in ∼20% cells death at 11 dpt^27^. To investigate whether the loss of USP19 affects the levels of pSyn in neuronal cells, we generated USP19 WT/M83^hem^ and KO/M83^hem^ primary mouse cortical neurons and performed IF imaging. As expected, PFF treatment led to a time-dependent pSyn accumulation in both USP19 WT and KO neurons. However, there was ∼50% reduction in pSyn fibrils at 10 dpt in KO neurons when compared to WT (Fig. 5A, B). To verify that this effect was not due to differences in PFF internalization between WT and KO neurons, we performed live- and fixed-cell time course measurements of uptake of fluorescently labelled PFFs. We observed that uptake begins at 1h of PFF exposure. PFFs are then gradually trafficked into LAMP1-positive lysosomes. We found no difference in the rates of internalization of PFFs between WT and KO neurons. (Fig. S4 and S5A, B). We next used cell viability assay to assess differences in PFF-induced cell death between WT and KO cells which could underlie the effect on pSyn accumulation. PFF treatment led to ∼20% cell death at 10 dpt in both USP19 WT and KO neurons (Fig. 5C). No differences in cell viability were observed in the absence of exposure to PFF. We also tested whether the loss of USP19 affected viability during neuronal differentiation and maturation in culture which could predispose KO neurons to lower pSyn accumulation. We did not observe any differences in maturation viability between genotypes (Fig. S5C).

**Fig. 5:**
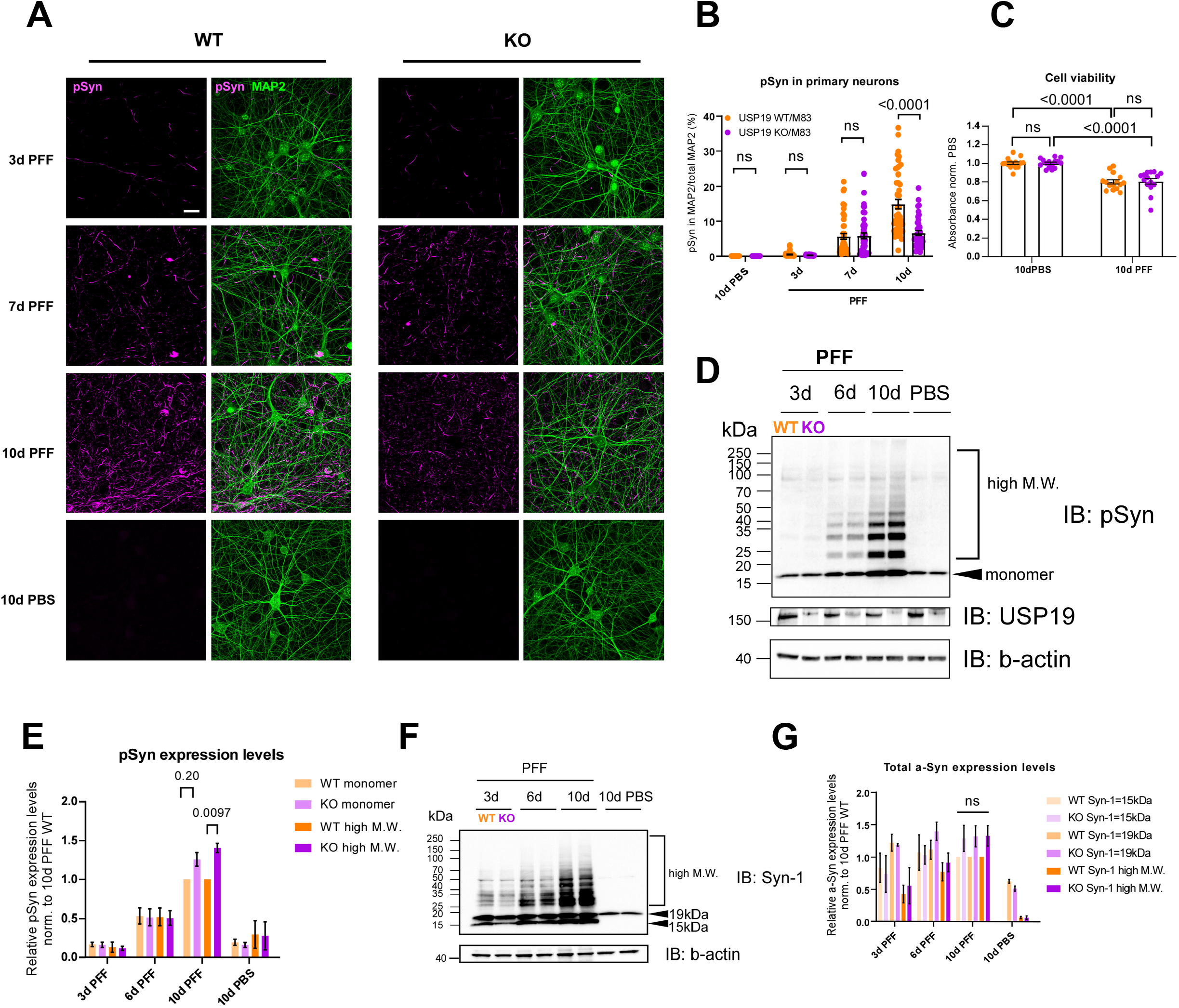
Loss of USP19 reduces pSyn pathology in primary mouse neurons. **A**. Representative confocal images of primary neurons (at 19 days in vitro [19 DiV]) treated with PFF or PBS for 3, 6 and 10 days and immunofluorescently labeled with MAP2 (neuronal marker) and pS129-Syn. Scale bar, 20 μm. **B**. Quantification of pS129-Syn and MAP2 cooccurrence represented as percentage area of pS129-Syn over total MAP2 area. 2-3 technical replicates (coverslips) and 5 field images per coverslip were acquired per culture. Each data point represents the percentage value per image. n= 3 biologically independent cultures. Two-way ANOVA followed by Tukey’s multiple comparisons test was used for statistical analysis. **C**. Quantification of cell viability test upon 10-day PBS or PFF treatment of primary neurons. n= 5 biologically independent cultures. Two-way ANOVA followed by Tukey’s multiple comparison test was used for statistical analysis. **D**. Representative immunoblots of total soluble pSyn in primary cortical neurons exposed to PFF or PBS for indicated number of days. **E**. Quantification of monomeric and high M.W. pSyn species. n= 5 biologically independent cultures. Two-way ANOVA followed by Tukey’s multiple comparison test was used for statistical analysis. **F**. Representative immunoblots of total soluble a-syn in primary cortical neurons exposed to PFF or PBS for indicated number of days. **G**. Quantification of fragmented and monomeric (15 and 19 kDa, respectively) and high M.W. a-Syn species. n= 3 biologically independent cultures. Two-way ANOVA followed by Tukey’s multiple comparison test was used for statistical analysis.

We next used immunoblot analysis to assess a-syn and pSyn species in WT and KO cells exposed to PFF. Cell lysates were subjected to centrifugation to try to isolate insoluble aggregates, but none were detectable probably due to the minimal amount of insoluble protein available from these primary neuronal cultures. In the *soluble* fraction, we did see a time-dependent increase in both monomeric and high molecular weight (M.W. bands >19 – 150 kDa) pSyn species in both WT and KO neurons. At 10 dpt, there was a small, but statistically significant increase in higher molecular weight species of pSyn in KO compared to WT neurons, possibly due to multiple posttranslational modifications, oligomerization, fragmentation of aggregates or combinations of these (Fig. 5D, E). A similar tendency to an increase in total higher molecular weight a-syn species was measured in KO neurons (Fig. 5F, G). The increase in *soluble* high molecular weight species in KO neurons contrasts with the decrease in *insoluble* high molecular weight species in KO brain (Fig 2G) raising the possibility that USP19 regulates the dynamics between soluble and insoluble pSyn species. Interestingly, there was also a tendency to more total fragmented and monomeric a-syn (15 and 19 kDa, respectively) at 10 dpt in KO neurons.

To test whether USP19 is involved in neuron-to-neuron propagation of a-syn pathology in an in vitro system, we used a microfluidic device that compartmentalizes PFF-receiving and non-receiving neurons which are in contact solely via axons. We then observed the development of pSyn pathology in the non-PFF exposed WT and KO neurons. We did not observe any differences in pSyn development in the non-exposed neurons regardless of whether WT or KO neurons were exposed to PFFs, suggesting that neuronal USP19 is not involved in neuron-to-neuron propagation mechanism of a-syn (Fig. S6).

### The loss of USP19 increases ubiquitination of a-syn containing aggregates

Since Lewy body inclusions are known to be ubiquitinated, we tested whether loss of the USP19 deubiquitinase might increase ubiquitination of a-syn. We therefore co-stained sections of WT and USP19 KO brains from animals treated with a-syn PFF with antibodies against pSyn and ubiquitin (Ub). We showed that pSyn^+^ aggregates are more ubiquitinated in many regions of the brains of USP19 KO animals (Fig. 6A, B). To test whether USP19 regulates the ubiquitination levels of pSyn and a-syn, we subjected soluble lysates of WT and KO primary neurons at 10 dpt with PFF or PBS to immunoprecipitation with anti-Ub antibodies and blotted the pellets with anti-pSyn or anti-syn antibodies. Multiple discrete bands 25 kDa or larger as well as high molecular weight smears were seen in PFF treated neurons that were either not detectable or diminished in the PBS treated neurons (Fig 6C, D). Bands at ∼15 kDa, the size of monomeric a-syn, or smaller were also detected which may reflect monomers or fragments of a-syn that were in complex with ubiquitinated a-syn. Quantification revealed significantly higher levels of ubiquitinated a-syn and a strong tendency to an increase in ubiquitinated pSyn in KO neurons treated with PFF (Fig. 6C, D). Reverse immunoprecipitation with anti-pSyn and blotting of the pellets with anti-Ub antibody confirmed our results of significantly higher levels of ubiquitinated pSyn in the KO (Fig 6E). Control immunoblots confirmed that the immunoprecipitations were successful in pulling down ubiquitinated proteins and total pSyn (Fig. S5D). These results suggest that USP19 regulates the ubiquitination of p-Syn, but we cannot exclude the modulation of ubiquitination of other proteins including non-phosphorylated forms of a-syn.

**Fig 6:**
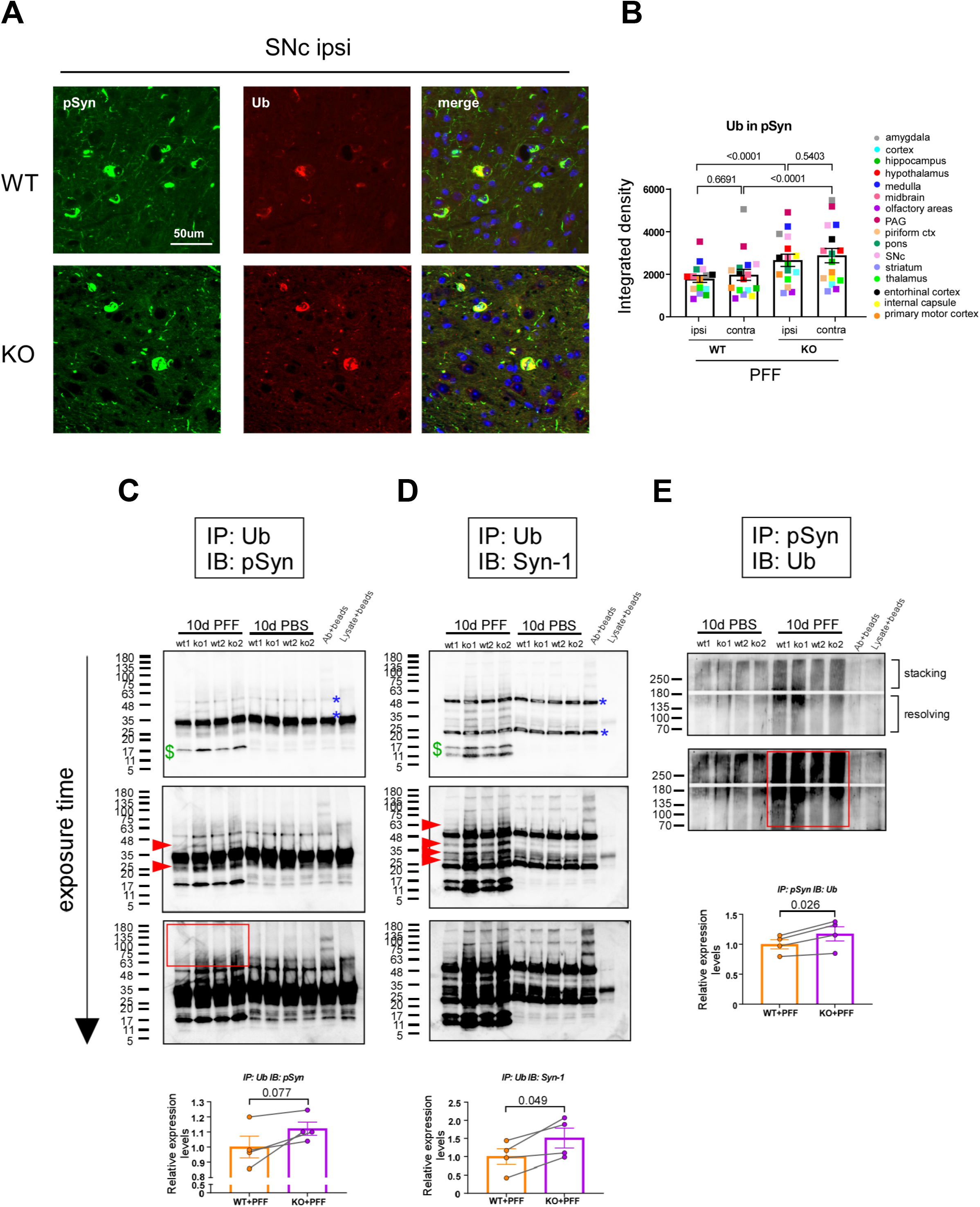
**A**. Representative IF images of pSyn and ubiquitin (Ub) colocalization in FFPE ipsi SNc. Sum of all pixel intensities (integrated density) of Ub was measured in pSyn-positive areas. **B**. Graph shows overall quantification of total brain Ub in pSyn areas of USP19 WT and KO/M83^hem^ animals at 60 dpi. n= 3 biologically independent animals. Each data point is a mean of means per brain region. Two-way ANOVA was used for statistical analysis followed by Tukey’s multiple comparisons test. P-values are indicated. **C-D**. Immunoprecipitation of ubiquitinated proteins from WT and KO neurons at 10dpt using anti-Ub antibody. IP samples (40% of total) were blotted for pSyn (H) and total a-syn (Syn-1, I). Graphs show whole lane quantification of PFF treated IP samples. Dollar symbols point to monomeric bands at the expected size of monomeric pSyn and a-syn. Arrowheads points to other specific Ub-pSyn and Ub-a-syn bands. Red rectangle shows high Ub-pSyn M.W. species. Blue asterisks point to heavy and light immunoglobulin chains. **E**. Reverse immunoprecipitation using anti-pSyn antibody. Both stacking and resolving gels were transferred and blotted for the presence of Ub-pSyn. Graph shows the quantification of high M.W. Ub-pSyn of PFF treated samples. n= 4 biologically independent cultures. Paired t-test was used for statistical analysis. Data are means ±s.e.m.. P-values are indicated.

## DISCUSSION

In this report, we demonstrate for the first time a role for the USP19 deubiquitinase in mediating the progression of Parkinson’s disease pathology in a mouse model. Using a murine model expressing the disease associated a-syn A53T mutation, we demonstrated that loss of USP19 clearly decreases the burden of pSyn positive aggregates in the brains injected with a-syn PFF. Other deubiquitinating enzymes such as USP30 and USP35 have been implicated in Parkinson’s disease through their roles in regulating mitophagy^58-60^. In addition, USP8 targets a-syn K63-linked polyubiquitin chains and is upregulated in human neurons with LB pathology. Interestingly, knocking down USP8 results in accelerated lysosomal degradation of a-syn and an improved rough-eye phenotype and locomotor function in a Drosophila model of a-syn induced toxicity^61^. On the contrary, USP9X is downregulated in SNc of PD patients and its deubiquitination of monoubiquitinated a-syn promotes proteasomal degradation^62^. These studies provide evidence of DUB’s differential regulation of a-syn stability. However, to our knowledge, this is the first demonstration of a role for a deubiquitinating enzyme in modulating specifically the accumulation of p-Syn containing aggregates which are the hallmark of PD. The improved pathology in the USP19 KO brains was evident in many regions and included substantia nigra pars compacta, striatum, hypothalamus, thalamus and cortex. These areas are all part of the cortico-basal ganglia-thalamo-cortical loop which controls sensorimotor function among others and its hypofunction is directly implicated in PD. Consistent with this, we observed a significantly better performance in the tail suspension test in the KO mice. In addition to the decrease in insoluble pSyn aggregates, we also observed a marked decrease in p-tau positive insoluble aggregates. This is in line with studies showing that a-syn promotes tau seeding and spreading in the context of PD and AD^27^.

We demonstrated here that USP19 was expressed in multiple cell types in the brain – neurons, astrocytes and microglia. The exact cell type(s) in which USP19 acts to support the accumulation of aggregates is unclear. However, we were able to demonstrate in primary neuronal cultures that loss of USP19 also decreases the accumulation of pSyn aggregates upon exposure to preformed fibrils indicating that USP19’s actions in neurons is at least partly responsible. Microglial activation in response to injected PFFs was also diminished significantly in USP19 KO brains (Fig. 3). This was revealed not only by decreased expression of markers such as Iba1 and GFAP, but also by the presence of smaller microglia. Since neuroinflammation has been implicated in the progression of PD (reviewed in^63^), it is possible that loss of USP19’s actions in these cells may also play a role in the neuroprotection seen in the KO mice. However, previous work in peripheral mononuclear cells has shown that loss of USP19 results in a hyperinflammatory response^27^. Thus, it is more likely that the decreased microglial activation in the KO brains was due to the decreased pathology causing less neuroinflammation.

USP19 KO decreased accumulation of pSyn aggregates and this was evident in the hemisphere injected with PFF and also in the contralateral hemisphere. This did not appear to be due to decreased uptake of the injected PFF as the number of cells affected by pSyn pathology in their cell bodies at an earlier time point when pSyn aggregates become detectable was similar in both WT and KO (Fig. 2D). In support of this, the rate of uptake of a-syn PFF was similar in WT and KO primary neuronal cells (Fig. S4 and S5A, B). Previous work on USP19 (ER isoform) indicated that it plays an important role in an unconventional pathway of protein secretion (MAPS) that appears specific for misfolded proteins including a-syn and tau^32^. We have measured levels of a-syn in the blood of these mice but did not detect any differences between WT and KO mice (data not shown). Supporting a lack of effect on secretion, we did not find any differences in the rate of propagation of pathogenic a-syn species between WT and KO cells using microfluidic chips that separate treated and non-treated neurons which connect solely by extending axon. (Fig. S6). We have attempted to measure the rate of secretion of a-syn from primary neurons derived from the WT M83 and USP19 KO M83 embryos and exposed to PFF. However, these measurements have been technically difficult due to the neurons not tolerating well the change of media required for such studies. The injection of PFF also induced the accumulation of p-tau aggregates and like the pSyn aggregates, their levels were decreased in the KO brains (Fig. 2H, I). Although the MAPS pathway can secrete not only a-syn but also tau and TDP-43 ^34^, it may not be the central mechanism regulating the levels of pSyn in our model.

In the absence of any effects of loss of USP19 on a-syn uptake or propagation, we conclude that USP19 regulates the intracellular economy of a-syn. In support of this, WT and KO mice examined at 60 days post injection of PFF had similar numbers of pSyn positive cells in multiple regions of the brain, but total pSyn pathology (area of pSyn) was reduced in the KO indicating a decrease in the amount of aggregates per neuron (Fig 2D, E). A similar decrease in pSyn aggregates was seen by IF in KO primary neuronal cultures chronically exposed to PFF (Fig 5A, B). This was coupled with an accumulation of soluble higher molecular weight species of p-Syn (Fig 5D, E) detected by immunoblotting, representing PTM modifications and/or oligomers of pSyn. Importantly, we found that pSyn^+^ aggregates were more ubiquitinated in the brains of USP19 KO animals (Fig. 6A, B). Immunoprecipitation experiments of soluble primary neuron cell lysates confirmed that USP19 KO have higher levels of ubiquitinated pSyn (Fig. 5H-J). These results provide evidence for USP19 regulation of ubiquitination of pSyn in both aggregates and soluble species. The cellular consequences of this altered ubiquitination are unclear at this time, but our results suggest at least two possible mechanisms (Fig 7). Increased ubiquitination of the soluble species may impair formation of aggregates as has been observed in vitro^64,65^ and increased ubiquitination of aggregates in the KO brain may result in lower levels of aggregates due to more efficient clearance in the KO. The lysosomal system is generally believed to play an important role in a-syn degradation^66-68^ and ubiquitination can target proteins to this system as well as to the proteasome. It is also possible that the increased ubiquitination promotes fragmentation or disassembly of aggregates into soluble oligomeric species. USP19 can interact with Hsp70 and this chaperone has been implicated in the disassembly of a-synuclein fibrils^69-71^.

**Fig. 7:**
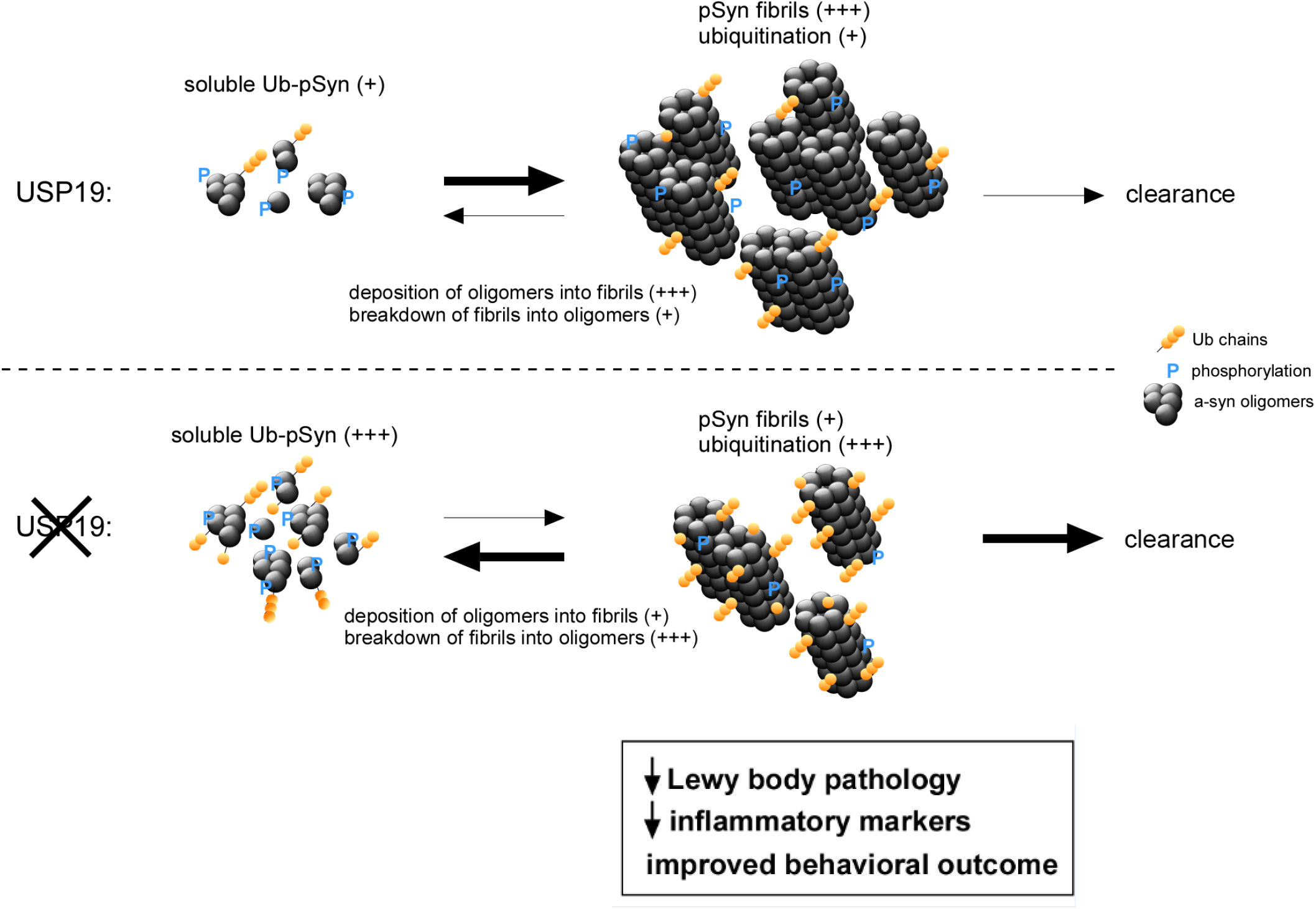
Roles of USP19 in modulating a-syn homeostasis. USP19 regulates the levels of ubiquitination of pSyn aggregates. Loss of USP19 does not affect a-synuclein uptake nor propagation but results in lowering of levels of aggregates due to impaired formation or enhanced clearance via the proteasomal or lysosomal systems and/or enhanced disassembly through USP19 regulation of chaperonin activities. The decreased Lewy body like aggregates results in lower levels of neuroinflammatory markers as well as improved behavioral outcome in USP19 KO.

Interestingly, we observed worse pSyn pathology in USP19 WT females compared to males (Fig. S2G, H). To our knowledge, this sex specific difference has not been previously reported in this commonly used model of PD and may reflect the fact that we used a large number of animals with both sexes in our study. The male to female prevalence of PD is about 3:2, although sex specific differences in severity of Lewy body pathology in humans have not been reported. Interestingly, male PD patients present with more rigidity, difficulties with vocal pitch, writing and gait and have higher prevalence of REM sleep disorder. On the other hand, females display more postural instability, freezing, are more likely to be depressed, and need higher levopoda dose (reviewed in^72^). The sex differences in pSyn pathology in our mice were not uniformly present in all brain regions. Such differences occurring in a regional manner could explain the sex-based differences in symptomatology described above. USP19 expression has been shown to be higher in female muscle due to the presence of an estrogen receptor α response element in the first intron of the USP19 gene^44^. Expression of this receptor has been reported to be 1.6x higher in female than male adult mouse brain^73^. Whether these findings can explain the sex specific differences we observed remains to be explored. Nonetheless, these observations reinforce the importance of conducting studies in adequate numbers of both sexes.

Although a clear decrease in pSyn pathology was evident in the KO brains, we did not observe an improvement in survival in the KO animals. One possible explanation for this phenotype might be unrelated to the Lewy body pathology. USP19 whole body KO animals have decreased body weight, mainly due to a significant reduction in the adipose tissue (^74^ and Fig. S3B). Thus, the reduced adipose tissue reserves in these global USP19 KO mice may have made them more prone to early death during PD-like disease progression. Future studies are needed to mitigate this confounding phenotype by knocking-out USP19 specifically in the CNS or even in specific subsets of neuronal or glial cells. The combination of PFF injection with endogenous production of mutant a-syn yielded a model in which disease progression was more rapid than what typically occurs in humans and so may also have obscured a survival benefit of loss of USP19. A clearer insight into this question is important in view of USP19 being a candidate gene for PD^40^ and as USP19 inhibitors have recently been described^75^ and might be a therapeutic approach for slowing the progression of PD or other misfolded protein associated neurodegenerative diseases.

## METHODS

List of antibodies used can be found in Supplementary Table 1.

### Animals

All animal studies were approved by the Animal Care Committee of the Research Institute of the McGill University Health Centre Research Institute. Animals were housed in high efficiency particulate air (HEPA)-filtered cages with a 12-hour light/dark cycle at 21°C with environmental enrichment and access to food (32% protein, 14% fat and 54% carbohydrates) and water ad libitum. The mice were generally housed in groups (maximum n=5 per cage), but males were frequently isolated due to aggressive behavior. M83 hemizygous (M83^hem^) mice expressing human alpha-synuclein with the A53T mutation under the control of a prion promoter (Pmp-SNCA*A53T;^41^) were obtained from The Jackson Laboratory. M83^+/+^ mice were mated with Usp19^−/−^/C57BL/6 mice (generated in our laboratory as previously described in^76^ for two generations to obtain M83^+/+^/USP19^+/-^. As M83^hem^ mice were used in this study, M83^+/+^/USP19^+/-^ mice were bred with USP19^+/-^ negative for M83 transgene to finally generate USP19^+/+^/M83^hem^ (USP19 WT/M83^hem^) and USP19^-/-^/M83^hem^ (USP19 KO/M83^hem^). All available animals (females and males) were used for each experiment; thus, groups were assigned based on genotype unless stated otherwise. Animals were genotyped using PCR-based amplification of M83 transgene and geo cassette present in USP19 deletion allele^76^.

### PFF preparation

Human a-syn PFF were prepared as previously described^54^. It was expressed in E. coli followed by endotoxin removal after purification. Monomeric a-syn (5 mg/ml) was allowed to spontaneously form fibrillar aggregates at 37°C in an orbital shaker for 5 days at 1,000 RPM. Quality control was assessed per batch using TEM to examine fibrillar morphology after negative staining, and Thioflavin-T assay to verify formation of amyloid fibrils, which were then sonicated by 40 cycles of 30 sec ON-OFF on a Bioruptor Pico (Denville, NJ) to yield fibrils of 50 to 100 nm long in hydrodynamic diameter as shown by dynamic light scattering^30,77^. Sonicated fibrils were aliquoted and stored at -80°C for short periods of time. An aliquot of PFFs was thawed at room temperature immediately prior to injection or culture treatment.

### Stereotaxic injection

Unilateral stereotaxic injection to deliver PFF or PBS into the right dorsal striatum was performed as previously described^42^. Briefly, at 3 months old, mice were positioned in the stereotaxic frame. A hole was drilled through the skull and injection performed at the following coordinates relative to bregma: A/P(y)= +0.2mm, M/L(x)= +2.0mm and D/V(z)= -2.6mm. Automatic injection (syringe: 5 µ l, 75 RN, 700 Series, Hamilton) was performed to administer the PFF (2.5 µ l of 5 mg/ml) at a rate 0.25 µ l/min. The wound was sutured, and animals were monitored closely until full recovery. Unless otherwise indicated, mice were sacrificed at 3 months post-injection.

### Histology

Mice were anaesthetized and transcardially perfused with ice-cold PBS followed by 4% paraformaldehyde (PFA) buffered in PBS. Brains were removed and post-fixed with 4% PFA in PBS at 4°C O/N. Brains were coronally pre-sectioned into 6 equally thick rostro-caudal parts, processed (dehydration and wax infiltration) and paraffin embedded. Six-micron sections were prepared with microtome (Leica), mounted onto microscopic slides, dried at 37°C and stored at RT until use.

#### Immunohistochemistry

Brain tissue on glass slides was deparaffinized and re-hydrated as follows: 3x 5min CitriSolv, 2x 5min 100% ethanol, 2x 5 min 95% ethanol, 1x 5min 70% ethanol and 2x 1 min deionized water. For antigen retrieval, slides were submerged in sodium citrate buffer (10 mM, pH=6) and boiled for 10 min. After cooling to RT, the slides were washed with 1X Tris-buffered-saline-TritonX10 (TBST, pH=7.6). Endogenous peroxidase activity was quenched for 15 min in 3% H_2_O_2_ in H_2_O followed by 2x TBST washes. Sections were blocked in 5% goat serum in TBST for 1 hour at RT and incubated overnight at 4°C in a humidified chamber with primary antibodies. After 3x 5 min TBST washes, sections were incubated for 1 hour at RT with HRP-conjugated secondary antibodies followed by 3,3′-diaminobenzidine (DAB) incubation. Chromogen development was closely monitored, and reaction stopped in water. Hematoxylin was used as a counterstain; sections were then sequentially dehydrated using ethanol solution gradient (70%, 95% and 100%) and mounted using organic mounting media (Permount, Fisher Scientific). Whole-slide images were acquired using the Leica Aperio AT Turbo digital pathology scanner (objective: 20x/0.75 Plan Apo). Image analysis was performed using QuPath digital pathology software^78^.

#### Immunofluorescence

Brain tissue was processed as described above up until primary antibody incubation. Previously validated primary antibodies were used for immunofluorescence followed by secondary fluorophore-conjugated antibodies. DAPI was used to stain nuclei. Dehydration step was omitted, and sections were mounted using aqueous mounting media. Whole section images were acquired using the Zeiss AxioScan Z1 slide scanning system (10X objective). Image analysis was performed in QuPath digital pathology software^78^ with built-in ImageJ extension and home-made macros.

#### RNA in-situ hybridization

RNA in-situ hybridization was performed according to the manufacturer’s protocol RNAscope® Multiplex Fluorescent Reagent Kit v2 Assay. Briefly, formalin-fixed paraffin-embedded brain sections were deparaffinized by sequential incubation in CitriSolv (3x 5 min) and 100% ethanol (2x min). Slides were dried for 5 min at 60°C. Sections were incubated in RNAscope Hydrogen Peroxide for 10 min at room temperature. Target retrieval was performed in a target retrieval buffer using a steamer at 99°C for 15 min. Sections were rinsed in distilled water and cooled down to room temperature, then incubated in 100% ethanol and dried. RNAscope® Protease Plus buffer was applied and slides incubated for 15 min in RNAscope humidified oven at 40°C then rinsed with water. Specific probes [USP19-ER (ACD #847951-C2), USP19-cyt (ACD #847961-C3) or USP19 common (ACD#847941)] were then hybridized for 2h at 40°C, then washed 2x 2 min in a wash buffer. Sequential hybridization of amplification probes was performed. Hybridization of HRP probes was performed thereafter. Finally, hybridization of detection probes was performed using fluorescent Opal dies. Immunofluorescence for neuronal and glial markers was performed thereafter.

### Behavioral phenotyping

For all behavioral evaluation, the experimenter was blinded to treatment and genotype groups.

#### Survival

PFF- and PBS-injected mice were subjected to a survival study to assess differences in overt pathology between USP19 WT and KO/M83^hem^ animals. Post-surgery, mice were closely monitored (twice a week) for the presence of PD-like symptoms (decreased movement, tremors and unbalanced gait) and drop in body weight. Closer to the 3-month post-PFF timepoint, mice were monitored daily. If mice exerted overt deficits or lost >20% of initial body weight due to inability to feed and drink, they were considered to have reached the clinical endpoint and were euthanized. Survival data were analyzed using the Kaplan-Meier survival curve.

#### Tail-suspension test

Tail-suspension test (TST) was performed to assess behavioral changes reminiscent of PD pathology. A mouse was securely fastened with a medical tape by its tail and hung head-down in obstacle-free air. Mobility was recorded using ANYmaze (Stoelting Co.) automatic movement tracking software during a single 5-min test period.

#### Grip strength

Forelimbs and hindlimb strength were measured using a Grip Strength Meter (Columbus Instruments, Columbus, OH, USA). Habituation (4 trials per each leg pair) to the apparatus was performed one day before the test. Average of 4 consecutive trials was used for analysis on the test day. Three different time points (pre-injection [time 0], 60 and 90 dpi) were assessed.

#### Wire hang

The wire hang test of motor function was conducted using a modified protocol previously described^43^. A mouse was placed on the top of a standard wire cage lid. The lid was lightly shaken to cause the mouse to grip to the wires and then turned upside down onto a 40 cm tall empty box. The latency to fall off the wire grid was measured. A trial was stopped if a mouse remained on the lid longer than 15 min. Three different time points (pre-injection [time 0], 60 and 90 dpi) were assessed. The animals were not subjected to prior habituation.

#### Rota-Rod

To assess motor learning, coordination, and balance, mice were subjected to the Rota-Rod test (Stoelting Co., Ugo Basile) as previously done in^43^ with minor modifications. Briefly, each mouse was given a training session (three 5-min trials, 5 min apart) to acclimate them to the apparatus. During the test period (2 hr later), each mouse was placed on the rotarod with increasing speed, from 4 rpm to 40 rpm for the period of 5 min. The latency to fall off the rotarod was recorded. Each mouse received two consecutive trials and the mean latency to fall was used for analysis. Three different time points (pre-injection [time 0], 60 and 90 dpi) were assessed.

### Protein biochemistry

#### Sequential solubilization of total brain protein

Ipsi- and contralateral hemispheres (or right and left in non-injected controls) were processed using previously established protocols^79-81^ with the following adjustments. Tissue was homogenized at 20,000 RMP (Polytron) in 10 volumes (w/v) of 1X TBS (Tris buffered saline) buffer (containing protease and phosphatase inhibitor cocktail (Sigma, USA) and PR-619 (Life Sensors, USA), a broad deubiquitinase inhibitor at 4°C. Homogenate was centrifuged at 1,000 x g to remove cell debris. Supernatant (crude homogenate) was placed in a polycarbonate centrifuge tube and centrifuged at 100,000 x g for 1 hour at 4°C. Resulting supernatant was retained as TBS-soluble fraction. Pellet was resuspended by sonication (10s, 20 kHz, 40% sonicator) in 5 volumes of 1X TBS buffer containing 5% SDS at RT. Second round of ultracentrifugation was performed at 100,000x g for 30 min at RT. Supernatant was retained as SDS-soluble fraction. Finally, the pellet was solubilized in about 50-100 µ l of 1X TBS buffer containing 5% SDS and 8M urea by water-bath sonication for 10 min to attain complete protein solubilization. Resulting fraction is termed the Urea-soluble fraction.

#### WB

Brain samples were subjected to SDS-PAGE using gradient gels (5-20%). Transfer was performed in 25 mM Tris, 192 mM glycine, 20% methanol for 2 hours at a constant current of 350-400mA at 4°C. When different forms of a-syn were detected, membranes were washed for 5 min with PBS and incubated in a fixing buffer (4% PFA, 0.01% glutardehyde in PBS) for 30 min. After incubation, membrane was washed 3x 5 min in 1X TBST buffer and blocked with 5% skimmed milk in 1X TBST for 30 min - 1 hour (if phosphorylated forms of proteins were detected, 50mM NaF was added to blocking buffer).

#### Immunoprecipitation

Proteins lysates from cultured neurons were sonicated at 4 °C for 10s and clarified at 10,000 x g for 10 min. Then, proteins from supernatant were solubilized for 1 h at 4 °C under gentle rotation in lysis buffer (10 mM Tris-HCl, pH 7.5, 10 mM EDTA, 150 mM NaCl, 1% Triton X-100, 0.1% SDS) supplemented with a protease inhibitor cocktail, phosphatase inhibitor cocktail, proteasome inhibitor MG132 and deubiquitination inhibitor PR-619. 800 µ g of proteins were incubated with 3 µ g of mouse monoclonal anti-ubiquitin antibody (clone FK2) or 3 µ g of rabbit monoclonal anti-phophoSer129 a-syn (or their corresponding IgGs as IP control) for 3h at 4 °C and then overnight at 4 °C with 60 µ l of protein G-sepharose beads. Precipitates were washed three times with 1 mL lysis buffer and proteins were eluted by boiling the beads 5 min in βME-reducing sample buffer before SDS-PAGE.

### Primary mouse cortical cultures

Cortical neurons were prepared from embryos (E15-16) of pregnant M83^+/+^ USP19^+/-^ females (mated with USP19^+/-^ males negative for M83 transgene) as previously described^82,83^ with some modifications. Briefly, embryonic brains were kept in hibernation media (Hibernate-E, Invitrogen) supplemented with 2% NeuroCult™ SM1 (Cederlane) and 0.5 mM L-GlutaMAX, Gibco) for several hours at 4°C. Single embryo genotyping was performed to identify USP19 WT and KO embryos. Cortices were then microdissected in ice cold HBSS (Gibco, 14170-112) supplemented with 33 mM sucrose, 10 mM HEPES and 0.5 mg/ml penicillin/streptomycin/glutamine (Gibco, 10378-16). Tissue was trypsinized and triturated using P1000 pipette in supplemented neurobasal medium (2% NeuroCult™ SM1 and 0.5 mM L-GlutaMAX). Cells (100,000 cells on 12 mm diameter coverslip or 500,000 cells on 35 mm culture plate or 6-well plate) were plated on poly-D-lysine-precoated surfaces in supplemented neurobasal media. Neurons were treated with PFF or PBS at 9-11 days in vitro (DiV) for the duration of 3, 7 and 10 days.

#### Immunofluorescence

Neurons treated with PBS or PFF for 3-10 days were fixed using 4% PFA, 5% sucrose in PBS for 1h. To block free aldehyde groups, 50 mM NH4Cl in 1X PBS was applied for 10 min. Following several PBS washes, 20 min blocking (10% serum, 0.02% TritonX-100 in 1X PBS) was performed. Cells were then incubated O/N at 4°C in primary antibodies in 1X PBS/5% serum/0.02% Triton-X100, then washed 3x 5min in 1X PBS at RT. Secondary antibodies were applied in 1X PBS/5% serum/0.02% Triton-X100 for 1h at RT. Following 3x 5 min PBS washes, coverslips were mounted on microscopic slides.

#### Microfluidics

Microfluidic chips (Omega, open top – eNuvio, Canada) were handled according to manufacturer recommendations. Briefly, chambers were coated with 0.1 mg/ml poly-D-lysine O/N. 70,000 cells were plated in each chamber. Fluidic separation of chamber #1 from #2 was maintained using hydrostatic pressure (higher media volume in chamber #2). PFF at 2 µ g/ml were used in the cell-to-cell propagation experiment. Immunofluorescence to label pSyn and MAP2 (dendritic marker) was performed following 10-day post-PFF treatment as described above. Microfluidic chips were imaged using Zeiss LSM780 confocal system with 10X objective, z-stacking and tile configuration.

#### Live-cell and fixed-cell imaging of PFF uptake

WT and KO neurons (16-18 DiV) were transduced with AAV to express GFP (3-4 days expression). Live-cell imaging was performed using 22 mm coverslips in a steel cell chamber (Attofluor™, Thermo Fisher). Neurons were kept on a heated stage (37 °C) on a Zeiss LSM780 inverted microscope. GFP and 568 fluorescence were excited through a x63 oil-immersion lens (NA, 1.4) using a 488-nm (1%) and 543-nm (3%) laser light, respectively. Time series (acquisition every 2 min) were collected for 1h as multiple image slice using the Zen software. In the fixed cell experiment, mature spiny neurons (18-21 DiV) were treated with 568-PFF (1μg/ml) for 1-24h, and IF was performed as described above. Multiple slice images (0.5 µ m range) were acquired on the Zeiss LSM880 microscope using x63 oil-immersion lens (NS, 1.4). The amount of internalized PFF was measured as mean fluorescence intensity in the cell body cytoplasm (avoiding membrane-attached PFF signal) using the ImageJ software.

### Image analysis

Histological images stained for pSyn were quantified using QuPath software and built-in Positive cell detection analysis. Brain regions were annotated, and number of total (hematoxylin) and positive cells (DAB) calculated using custom parameters. Each sample was then inspected for the presence of false detections (due to tissue, staining or imaging artefacts) which were then removed from analysis. Sections of similar Bregma positions were used for pSyn pathology comparison between groups. Four sections per animal were used for image analysis. Experimenter was blinded to the treatment and genotype.

## DATA AVAILABILITY

The authors declare that all data supporting the findings of this study are available within the paper and its supplementary information file. Any ‘data not shown’ is available from the corresponding author upon request.

## ACKNOWLEDGMENTS

This work was supported by the United States Dept of Defense Congressionally Directed Medical Research Programs of grant #PD170110 and by a Parkinson Canada Pilot Project Grant awarded to SSW, TD, and YY and Canadian Institutes of Health Research grant # FRN 168937 and Natural Sciences and Engineering Research Council Discovery Grant # RGPIN-2016-04054 to SSW. Y.Y. is supported by an intramural research program of NIDDK in the National Institutes of Health. LS and JBP were recipients of fellowship awards from the Research Institute of the McGill University Health Centre. HH was the recipient of a studentship award from the McGill Healthy Brains, Healthy Lives Canada First Research Excellence Fund and JH was the recipient of a Canada Graduates Scholarship from the Canadian Institutes of Health Research.

## COMPETING INTERESTS

SSW has performed contract research for Almac Discovery for studies not involving the results presented here.

## AUTHOR CONTRIBUTIONS

LS performed all histology, immunostaining, primary culture and imaging experiments. LS, NB, JBP and YW performed behavioural tests. LS and MPi performed image analysis. LS, NB and AKh performed biochemical experiments. LS and HH prepared primary neuronal cultures. LS and JH performed cell viability assay. WL prepared and EC and IS validated the PFF preparations. MP participated with animal breeding and colony maintenance. LS, SW, TD and YY contributed to hypothesis development, experimental design and data interpretation. SW provided overall supervision. SW, TD and YY provided funding. All authors discussed the data and commented on the manuscript.

---

**TABLE 1:** List of antibodies.

| Antibody | Source | Cat # | Dilution |
| --- | --- | --- | --- |
| NeuN | Millipore,<br>Abcam | MAB377,<br>Ab177487 | IHC-1:500 |
| Tyrosine<br>Hydroxylase | Millipore | AB512 | IHC-1:500 |
| GFAP | Millipore,<br>Abcam | MAB360,<br>Ab7260 | IHC-1:500 |
| Iba1 | Wako | 019-19741(IF) | IHC-1:500 |
| pSyn | Abcam | ab51253,<br>ab184674 | IHC-1:500<br>WB- 1:1000 |
| a-syn | BD Biosciences | 610787 | WB-1:750 |
| ptau | Thermo fisher | MN1020 | IHC-1:250 |
| synaptotagmin-1 | Synaptic systems | 105011 | WB- 1:1000 |
| SNAP25 | Synaptic systems | 111011 | WB- 1:1500 |
| MAP2 | Synaptic systems | 188044 | IF- 1:1000 |
| ubiquitin | Millipore | MAB1510<br>(Ubi-1) | IHC- 1:500 |
| ubiquitin | Millipore | 04-263 (FK2) | IP |
| b-actin | Millipore | A1978 | IB-1:8000 |
| USP19 | Abcam | Ab167059 | WB- 1:1000 |

