## Supplementary material for "Genetic inactivation of the USP19 deubiquitinase regulates a-synuclein ubiquitination and inhibits accumulation of Lewy body like aggregates in mice": (Fig. S1A, B)

### Supplementary figure 1

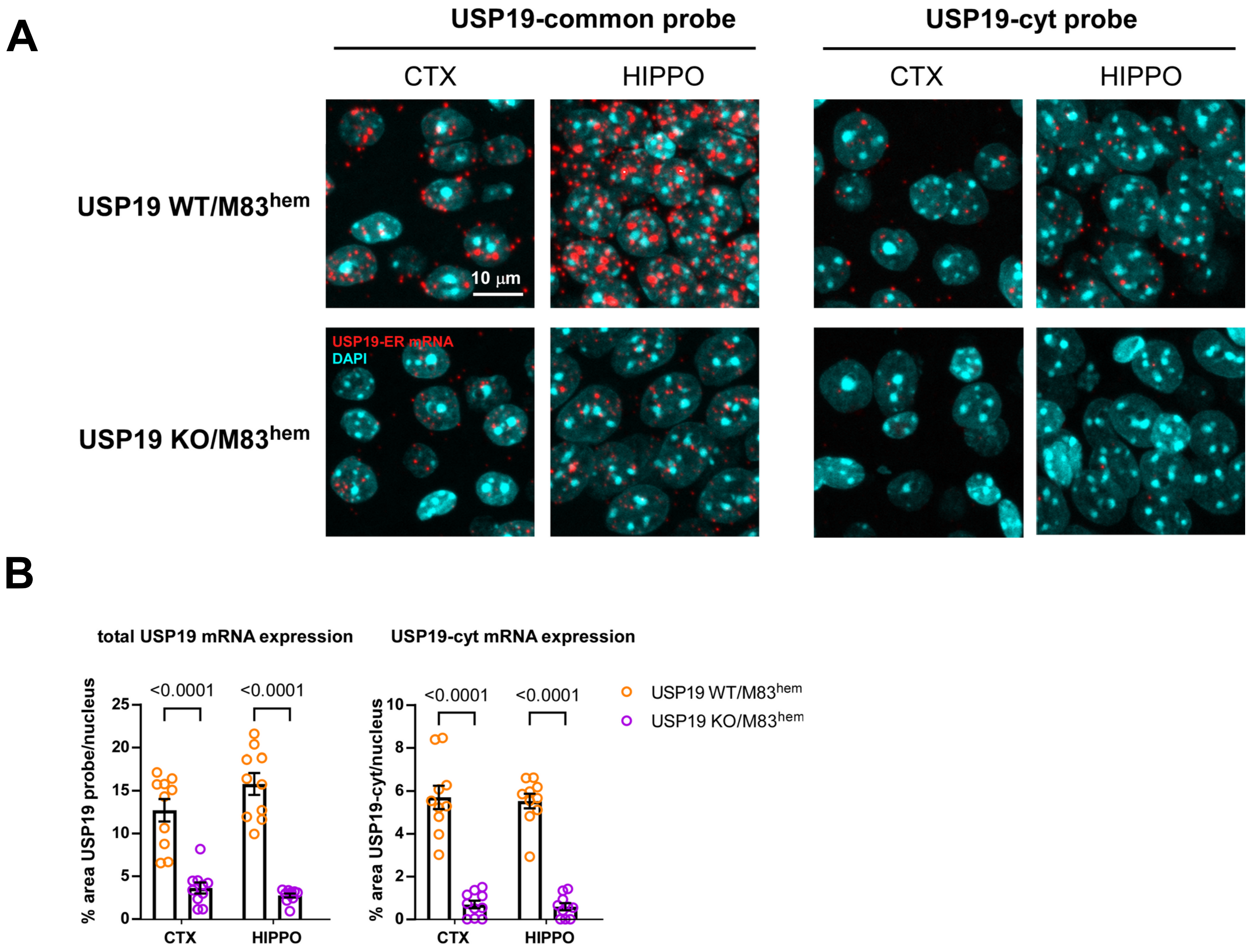


**Supp. Fig. 1: USP19 cytoplasmic isoform is expressed in the brain.**
**A.** RNAScope in-situ hybridization on FFPE brain sections to detect the presence (red signal) of both ER and cytoplasmic USP19 (using common probes) and cytoplasmic only USP19 isoform in USP19 WT/M83hem and USP19 KO/M83hem animals. Shown are representative images of CTX and HIPPO. DAPI was used to label nuclei. **B.** Quantification of RNAScope USP19-common and -cyt signals presented as percent area of red pixels in 10 randomly selected nuclei in CTX and HIPPO.

### Supplementary figure 2

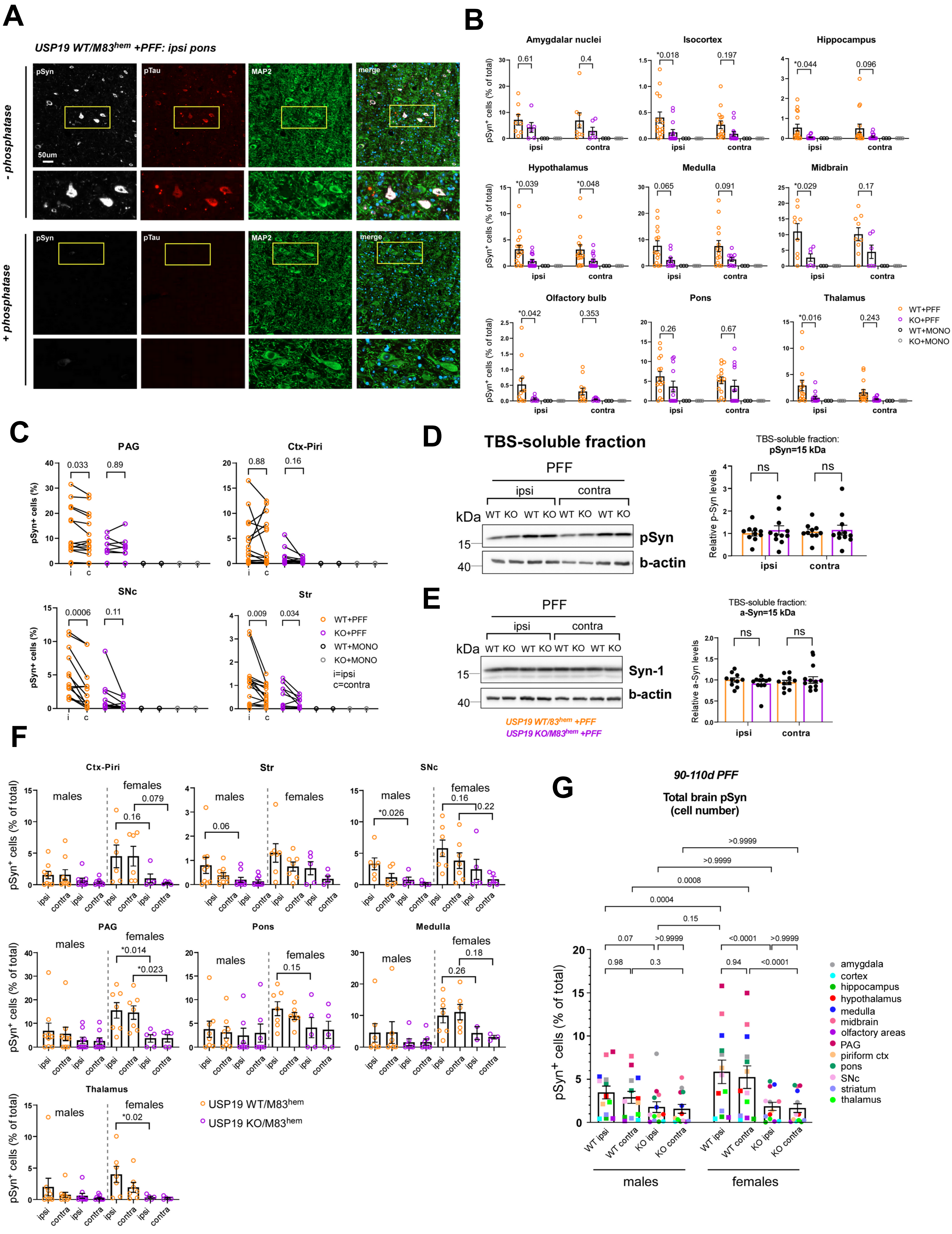

**Supp. Fig. 2: Levels of total soluble pSyn and total soluble a-Syn in USP19-WT and -KO PD-like brains. A.** Immunofluorescence images of pSyn (A) and pTau (B), and neuronal (MAP2) and DAPI staining of brain sections (shown is ipsi pons) treated or not with phosphatase to test the specificity of pSyn and pTau staining. **B.** Quantification of $pS129\text{-}Syn^+$ cells using QuPath analysis in indicated brain regions. Each data point represents a mean of 4 brain sections per animal. $n \geq 10$ (PFF-injected) and n=3-4 (a-syn monomers-injected) biologically independent animals. Data are mean ± s.e.m. Two-way ANOVA was used for statistical analysis followed by Sidak multiple comparisons test. **C.** Paired analysis of $pS129\text{-}Syn^+$ cells in indicated brain regions showing ipsi- to contralateral region spreading of pSyn pathology. Each data point represents a mean of 4 brain sections per animal. $n \geq 10$ (PFF-injected) and n=3-4 (a-syn monomers-injected) biologically independent animals. Data are mean ± s.e.m. Paired t-test was used for statistical analysis. **D.-E.** Representative immunoblots of pS129-Syn and total a-syn (Syn-1) of TBS-soluble whole hemisphere fractions. Beta-actin was used as a loading control. Data are mean ±s.e.m., $n \geq 10$ biologically independent animals. **F.** Quantification of $pS129\text{-}Syn^+$ cells using QuPath analysis in indicated brain regions in male and female mice. Each data point represents a mean of 4 brain sections per animal. $n \geq 4$ PFF-injected biologically independent male or female mice, except for medulla region ($n \geq 2$). Data are mean ± s.e.m. Two-way ANOVA was used for statistical analyses followed by Sidak multiple comparisons test. P-values are indicated. **G.** Overall quantification of total brain pS129-Syn represented as $pSyn^+$ cells (% of total) using IHC in A with additional brain regions at 90-110 dpi. Each point is a mean of means per brain region (each brain region is color-coded). Data are mean ±s.e.m. Two-way ANOVA was used for statistical analysis followed by Tukey's multiple comparisons test. P-values are indicated.

### Supplementary figure 3

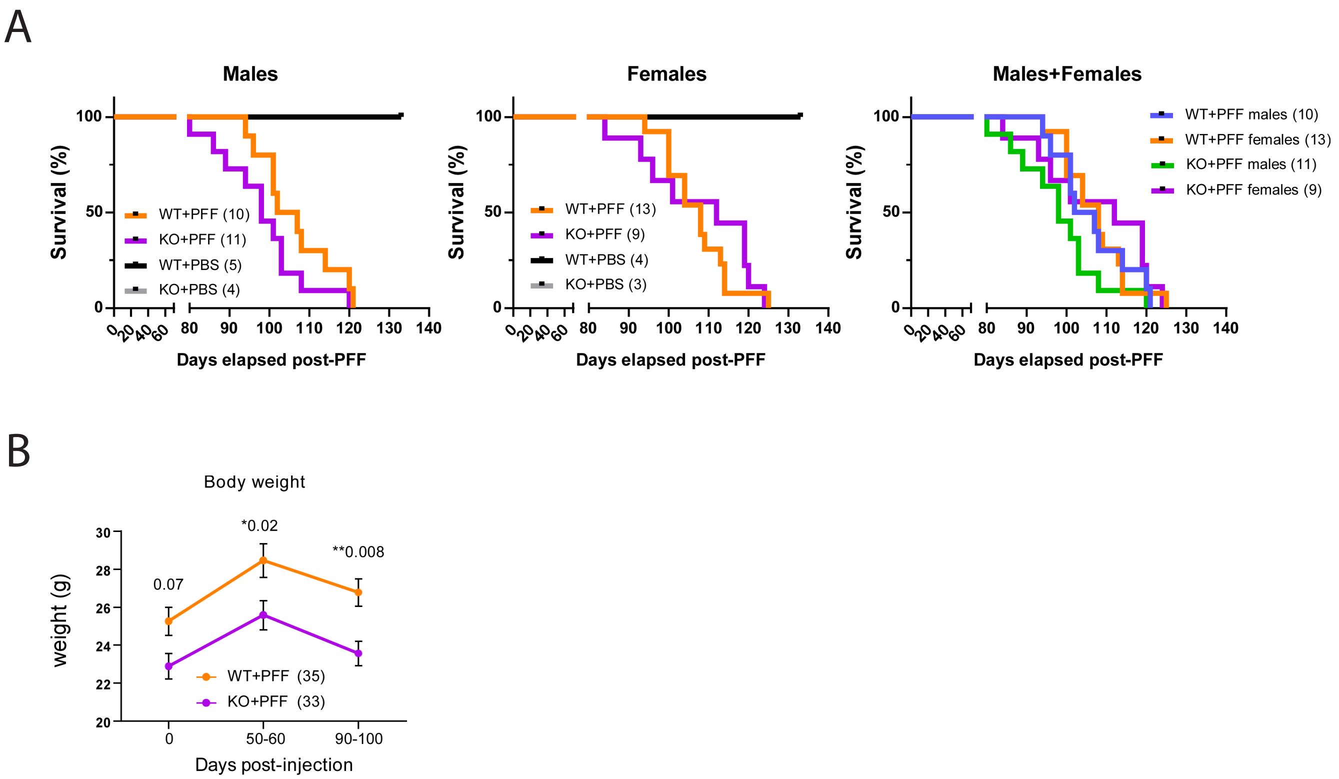


**Supp. Fig. 3: Survival of USP19 WT and KO PD-like mice**

**A.** Kaplan-Meier survival analysis of USP19 WT/M83hem and KO/M83hem males and females injected with PFF or PBS. n≥ 3 biologically independent animals. Statistical analysis for survival curves was performed by long-rank (Mantel-Cox) test. **B. E**ffects of USP19 depletion on body weight in PD-like mice. KO animals have significantly reduced body weight compared to WT which may lead to a faster deterioration upon the manifestation of PD-like symptoms.

### Supplementary figure 4

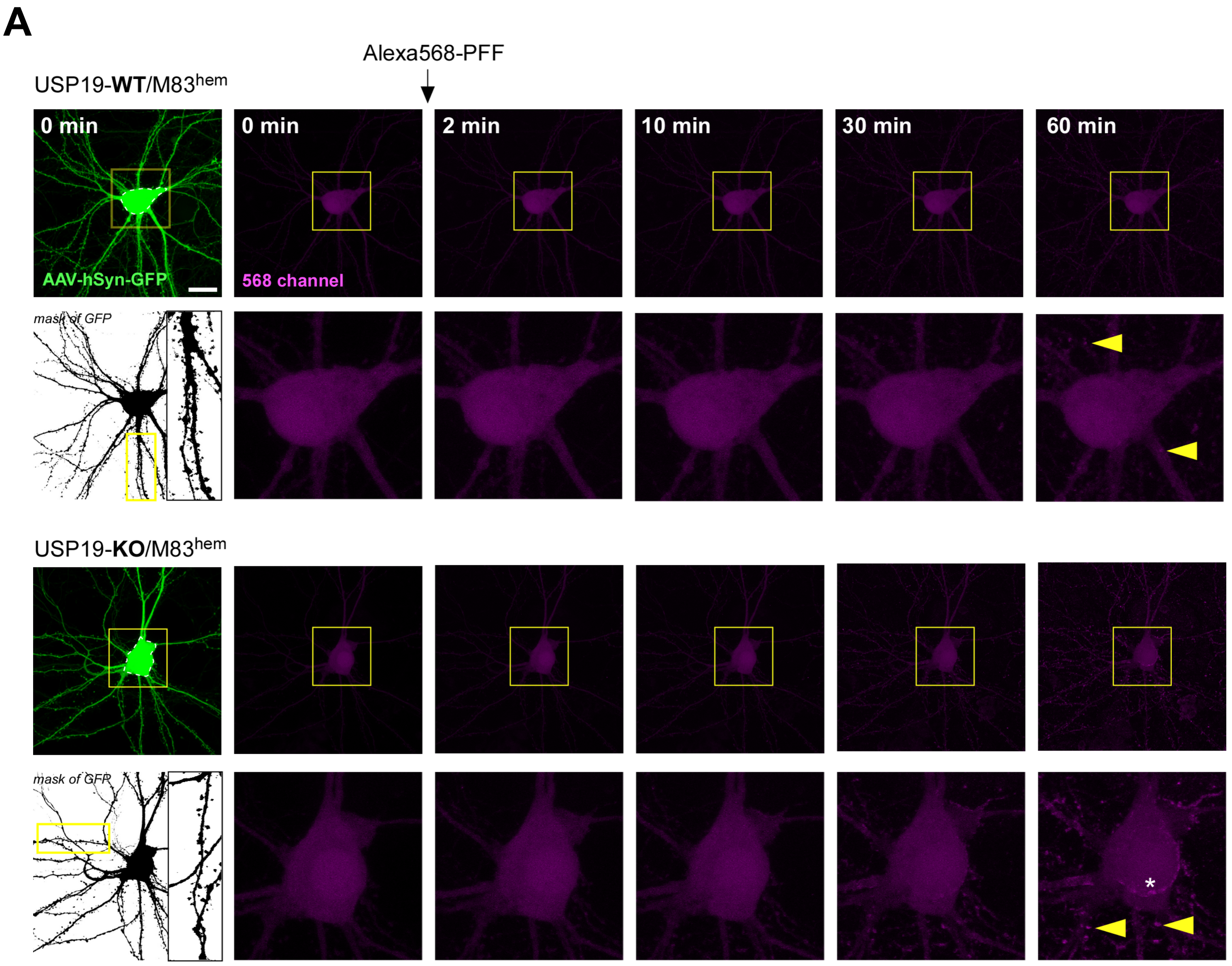


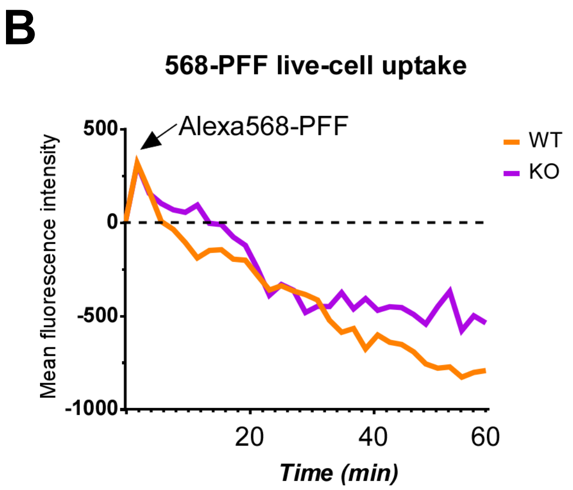


**Supp. Fig. 4: Live-cell PFF uptake in USP19 WT and KO neurons.**

**A.** Micrographs of live-cell imaging of Alexa568-PFF (magenta) uptake in WT and KO neurons. GFP-expressing mature (20 DiV) spiny neurons (B/W mask inserted for better visualization) were treated with 1 g/ml Alexa568-PFF and mean fluorescence intensity was measured in the cell body as measure of uptake over time. **B.** Quantification shows a decrease in mean fluorescence intensity in both WT and KO neurons over time. This is likely due to bleaching of the background fluorescence suggesting that 568-PFF uptake had not occurred under these conditions. Of note, asterisk shows that PFF are recruited to the membranes along the whole neuron (dendrites, spines and cell body). This was observed on similar time scales between WT and KO neurons (data not shown).

### Supplementary figure 5

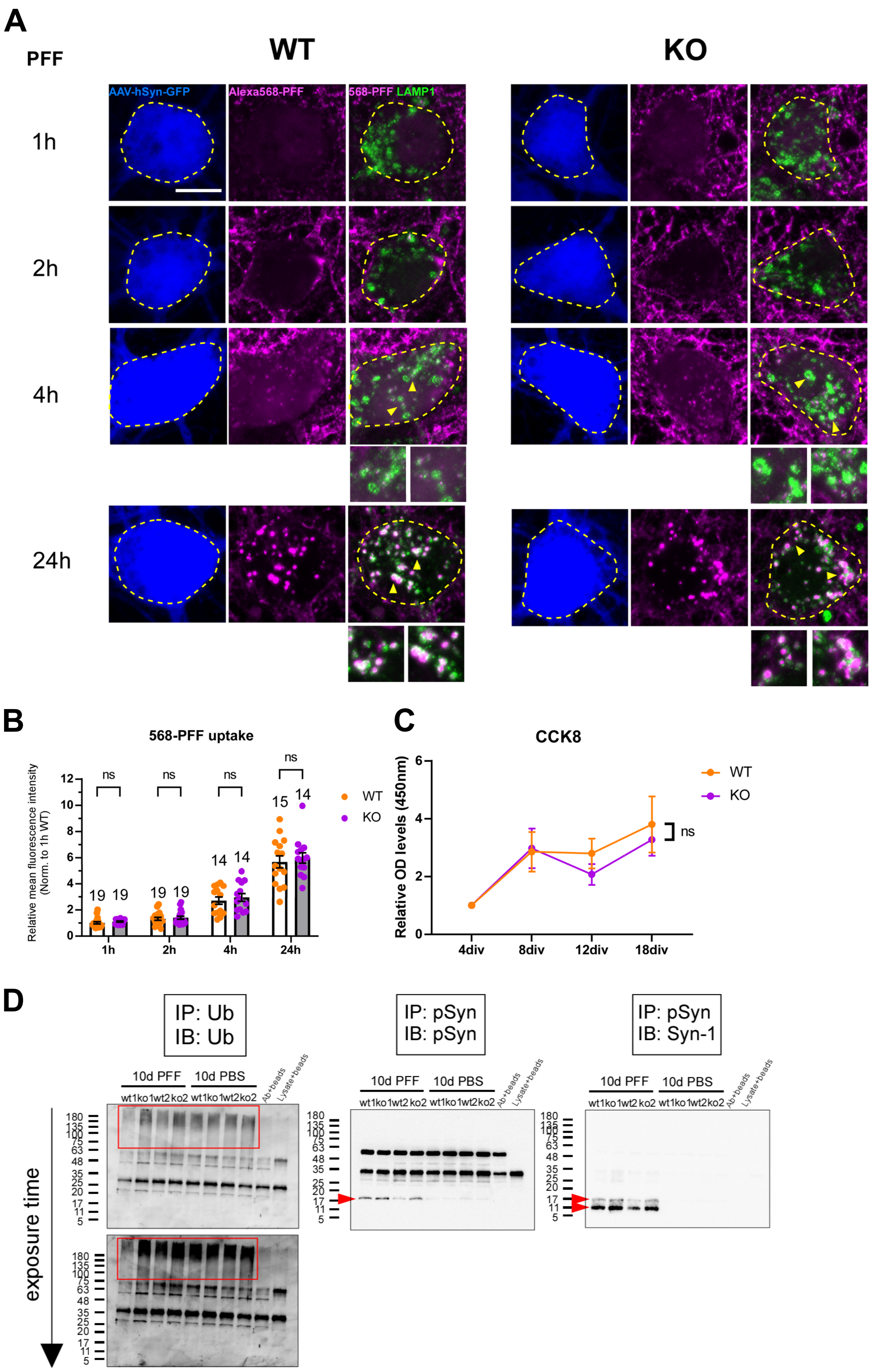

**Supp. Fig. 5: PFF uptake is indifferent between WT and KO neurons.**

**A.** Micrographs show a time course experiment of Alexa568-PFF (magenta) uptake in primary WT and KO neurons transduced to express GFP (blue) and immunolabeled for LAMP1 (green, lysosomal marker). Scale bar= 10 μm. **B.** Quantification shows mean red fluorescence intensity measured in the cell body over 24h post-PFF treatment. n= 3 independent cultures. Numbers above bars mark total number of neurons analyzed (>4 neurons per culture). Overall, very little uptake, if any, takes place in the first 2h post-PFF treatment. PFF puncta are clearly seen in colocalization with LAMP1 at the 4h time-point (higher magnification insets) as well as an increased diffused red signal in the cell body cytoplasm. At 24h, much of the membrane-recruited PFF (can be seen as strong signal around the cell body at previous timepoints) are now internalized and trafficked to lysosomes. **C.** Cell counting kit (CCK8) was used to assess differences in cell viability between USP19 WT and KO neurons during maturation in culture. Four time points (4, 8, 12 and 18 DiV) were assessed. n=3 independent primary cultures, each with three technical replicates (wells) per timepoint. Data are mean ±s.e.m. Two-way ANOVA followed by Tukey's multiple comparison test was used for statistical analysis. **D.** Immunoblot images show successful immunoprecipitation of ubiquitinated proteins and pSyn from WT and KO neurons at 10dpt using anti-Ub and anti-pSyn antibodies, respectively. IP samples (10% of total) were blotted for Ub, pSyn and total a-syn (Syn-1). Rectangles show high M.W. Ub-proteins. Arrowheads point to pulled down pSyn and a-syn.

### Supplementary figure 6

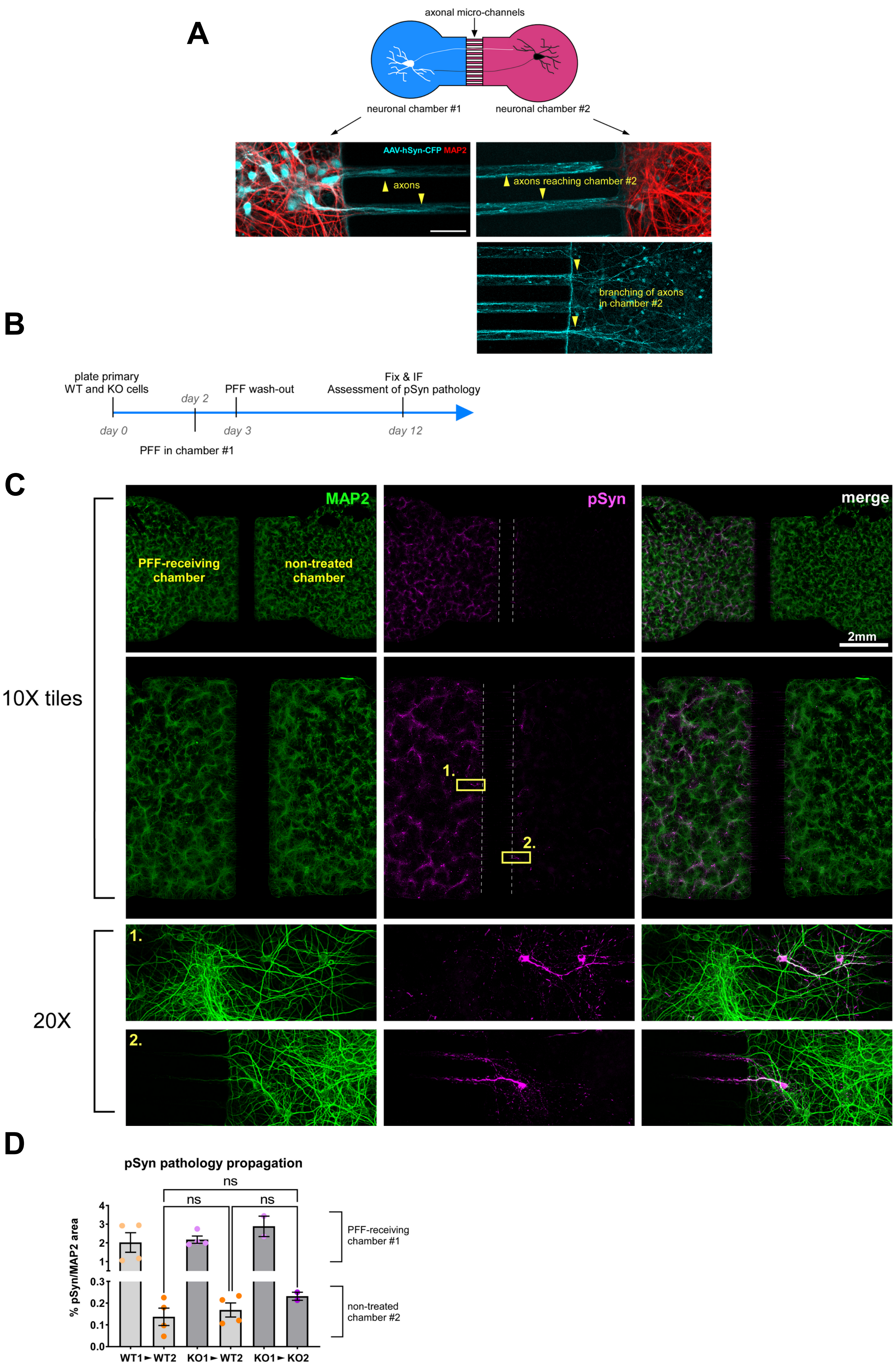

**Supp. Fig. 6: Loss of USP19 does not affect neuron-to-neuron propagation of pSyn pathology**

**A.** Scheme of a microfluidic device (Omega, Enuvio) composed of two neuronal chambers connected by axonal microchannels. Micrographs show good experimental conditions needed to study cell-to-cell propagation where axons of neurons in chamber #1 transduced to express GFP successfully reach to and branch out in chamber #2. **B.** Scheme of cell-to-cell propagation experiment. **C.** Micrographs of the cell-to-cell propagation experiment. Whole chamber tile images were acquired at 10X using confocal microscope. Insets at 20X show that neurons in both chambers #1 and #2 look healthy and develop $pSyn^{+}$ inclusions. **D.** Quantification of pSyn pathology in PFF-receiving and opposed chambers in WT to WT, WT to KO and KO to KO conditions. n= 2-4 independent cultures. One-way ANOVA was used for statistical analysis followed by Tukey's multiple comparisons test. Data are mean ±s.e.m.
